# DiatOmicBase, a gene-centered platform to mine functional omics data across diatom genomes

**DOI:** 10.1101/2024.09.12.612655

**Authors:** Emilie Villar, Nathanaël Zweig, Pierre Vincens, Helena Cruz de Carvalho, Carole Duchene, Shun Liu, Raphael Monteil, Richard G. Dorrell, Michele Fabris, Klaas Vandepoele, Chris Bowler, Angela Falciatore

**Author notes:** Department of Algal Development and Evolution, Max Planck Institute for Biology, 72076 Tuebingen, Germany (Carole Duchene).

## Abstract

Diatoms are prominent microalgae found in all aquatic environments. Over the last 20 years, thanks to the availability of genomic and genetic resources, diatom species such as *Phaeodactylum tricornutum* have emerged as valuable experimental model systems for exploring topics ranging from evolution to cell biology, (eco)physiology and biotechnology. Since the first genome sequencing in 2008, numerous genome-enabled datasets have been generated, based on RNA-Seq and proteomics, epigenomes, and ecotype variant analysis. Unfortunately, these resources, generated by various laboratories, are often in disparate formats and challenging to access and analyze. Here we present DiatOmicBase, a genome portal gathering comprehensive omics resources from *P. tricornutum* and two other diatoms to facilitate the exploration of dispersed public datasets and the design of new experiments based on the prior-art.

DiatOmicBase provides gene annotations, transcriptomic profiles and a genome browser with ecotype variants, histone and methylation marks, transposable elements, non-coding RNAs, and read densities from RNA-Seq experiments. We developed a semi-automatically updated transcriptomic module to explore both publicly available RNA-Seq experiments and users’ private datasets. Using gene-level expression data, users can perform exploratory data analysis, differential expression, pathway analysis, biclustering, and co-expression network analysis. Users can create heatmaps to visualize precomputed comparisons for selected gene subsets. Automatic access to other bioinformatic resources and tools for diatom comparative and functional genomics is also provided. Focusing on the resources currently centralized for *P. tricornutum*, we showcase several examples of how DiatOmicBase strengthens molecular research on diatoms, making these organisms accessible to a broad research community.

**Significance statement:** In recent years, diatoms have become the subject of increasing interest because of their ecological importance and their biotechnological potential for natural products such as pigments and polyunsaturated fatty acids. Here, we present an interactive web-based server that integrates public diatom ‘omics data (genomics, transcriptomics, epigenomics, proteomics, sequence variants) to connect individual diatom genes to broader-scale functional processes.

## Introduction

Diatoms are unicellular algae that play a major ecological role by contributing up to 20% of carbon fixation in aquatic ecosystems (Tréguer et al., 2021). The group encompasses up to 100,000 species (Alverson et al., 2007; Malviya et al., 2016) with a large diversity of morphologies, sizes and life histories. They are classified into two morphogroups according to their shape: centric diatoms are radially symmetric while pennates are bilaterally symmetrical. Their main common characteristic is the silica cell wall, or “frustule”, that make diatoms key players in the oceanic silica biogeochemical cycle (Tréguer et al., 2021) in addition to the carbon cycle. Widely distributed in any aquatic or humid environments, diatoms are particularly abundant in nutrient-rich coastal ecosystems as well as at high latitudes (Malviya et al., 2016). The diverse habitats in which they occur reflect their extreme adaptive capacities: they can be planktonic and benthic, and can be found even as epiphytes in terrestrial forests, in soil or associated with sea ice (Vanormelingen et al., 2009, Singer et al., 2021).

Besides their ecological significance, diatoms have recently emerged as interesting novel experimental systems to explore still largely uncharacterized features of phytoplankton biology (Falciatore and Mock, 2022). Two species are widely used to decipher diatom biology and molecular functioning: *Thalassiosira pseudonana* (a centric diatom) and *Phaeodactylum tricornutum* (a pennate diatom). They both present significant advantages: they are easily cultivated in the laboratory with rapid growth rates, their genetic transformation is mastered and they have small genomes (32.1 and 27.4 Mb, respectively) encoding around 12,000 genes (Armbrust et al., 2004; Bowler et al., 2008). In recent years, a growing number of different genomic, genetic, physiological and metabolic information and resources have been generated for these two species, making them suitable models to study diatom gene functions and metabolic pathways (Broddrick et al., 2019; Falciatore et al., 2020; Poulsen and Kröger, 2023). In parallel, additional diatom species have also been proposed as models to understand specific eco-physiological adaptations (*e.g*., *Fragilariopsis cylindrus* for polar habitats (Mock et al., 2017), *Thalassiosira oceanica* for open-oceans (Lommer et al., 2012), and *Seminavis robusta (Osuna-Cruz et al., 2020)* for benthic environments) or to address specific features of diatom life cycles (e.g., *Pseudo-nitzschia multistriata* for studying diatom sex (Ferrante et al., 2023)).

As of March 2024, 117 complete genomes of pennate and centric diatoms have been deposited in Genbank, and an ongoing project has announced the sequencing of 100 new diatom genomes (JGI initiative). The Marine Microbial Eukaryotic Transcriptome Sequencing Project (MMETSP) has additionally provided 92 different transcriptomes from diverse diatom species (Keeling et al., 2014) which has been recently completed with 6 other transcriptomes (Dorrell et al., 2021), providing a good representation of the most abundant diatom species in the ocean (Malviya et al., 2016). Finally, 54 single-cell and metagenome-assembled genomes (sMAGs) from diatoms have been assembled from meta-transcriptome/- genome data derived from *Tara* Oceans (Delmont et al., 2022), allowing complementary insights into the biology and ecology of uncultured and uncultivable species (Nef et al., 2022). Phylogenetic analyses of diatom genomes have revealed an extensive gene repertoire, which can be considered in a phylogenetic sense to constitute a patchwork coming from an ancient host, several endosymbionts acquired at different times, and bacterial horizontal gene transfers (Dorrell et al., 2021, Vancaester et al., 2020). Reflecting their complex evolutionary histories and phylogenetic distance (perhaps a billion years) from better-studied model eukaryotes within the animals, fungi and plants, the functional organization and evolutionary trajectories of diatom genomes are highly distinctive. These include families of novel transposable elements (Hermann et al., 2014), novel epigenomic marking of chromatin (Veluchamy et al., 2015; Zhao et al., 2021) and an apparent lack of structured centromeres (Bowler et al., 2008).

Combined functional genomic approaches in model species have been used to begin to decipher molecular actors regulating diatom physiology and distinct cellular and metabolic features. These include extensive energetic exchanges between plastids and mitochondria that augment CO_2_ assimilation (Bailleul et al., 2015), a peculiar organisation of the photosystems in the plastid membranes (Flori et al., 2017), and novel Light Harvesting Complex (LHC) protein families (Bailleul et al., 2010; Buck et al., 2019). The central role of diatoms in marine biogeochemical cycles has further been explored through identification of molecular mechanisms of nutrient uptake and metabolism, e.g., for silica (Nemoto et al., 2020), carbon (Shen et al., 2017), iron (Gao et al., 2021), nitrogen (Rogato et al., 2015) and phosphorus (Dell’Aquila and Maier, 2020). Comparative genomics and molecular physiology studies have greatly contributed to predict the role of nearly half of the diatom gene repertoire (Blaby-Haas and Merchant, 2019): as of July 2024, 6,781 genes from the nuclear genome (out of 12,357) have at least one annotation from Gene Ontology (GO), InterPro or Kyoto Encyclopedia of Genes and Genomes (KEGG) databases. Notwithstanding, the functional significance of many other diatom genes remains largely unexplored.

Among all studied diatom species, the genomic resources for *P. tricornutum* are the most advanced (Russo et al., 2023). The first assembly of the *P. tricornutum* nuclear genome constituted 27.4 Mb and 10,402 total gene models, whose annotation was assisted by the availability of an extensive collection of Expressed Sequence Tags (ESTs) (https://mycocosm.jgi.doe.gov/Phatr2/Phatr2.home.html) (Bowler et al., 2008). At the same time, a complete mitochondrial genome (77 kb, containing 60 genes (Oudot-Le Secq and Green, 2011)) and plastid genome (117 kb, containing 162 genes (Oudot-Le Secq et al., 2007)) were assembled. Following the initial assembly, the nuclear genome annotation was refined using 90 RNA-Seq datasets and more advanced annotation algorithms, resulting in the Phatr3 annotation. This annotation is available on the Ensembl archive (http://protists.ensembl.org/Phaeodactylum_tricornutum/Info/Index; Rastogi et al., 2018). More recent annotations (e.g., using proteomic data (Yang et al., 2018) and a new telomere-to-telomere *P. tricornutum* genome assembly (using long read sequencing technology (Filloramo et al., 2021 and Giguere et al., 2022)) have been performed, but to date no new annotation has been proposed to the community. Concerning transcriptomics, 123 microarrays have been used to generate a co-expression network (Ashworth et al., 2016), which is deposited on the DiatomPortal website (http://networks.systemsbiology.net/diatom-portal), while gene clustering of RNA-Seq experiments were recently used to create the PhaeoNet database (Ait-Mohamed et al., 2020). The epigenomic PhaeoEpiView browser including published epigenomic data has also been generated (Wu et al., 2023). Comparative genomics including *P. tricornutum* with a complete set of functional annotations are also provided in the PLAZA diatom comparative genomics platform (Vandepoele et al., 2013, Osuna-Cruz et al., 2020).

Despite the wealth of information generated from *P. tricornutum* summarized above, the resources are often disconnected, not always updated, or are incomplete, which diminishes their potential to yield valuable insights to the research community. To improve the connectivity between all the published omics data, we present here DiatOmicBase, a gene-centered web portal aiming to centralize all omics data related to diatoms. By focusing on the *P. tricornutum* model system, we provide different examples of the utility of this resource, e.g., from the analysis of gene and protein characteristics, as well as gene expression analyses under different conditions using previously published datasets. We demonstrate how querying the DiatOmicBase resources can enable a deeper understanding of diatom gene products and their roles in specific physiological and cellular processes, and whole cell metabolic fluxes that may be exploited for biotechnological and synthetic biology applications (Kumar et al., 2022). In addition to exploring existing datasets, the portal also allows submission of a user’s own data to provide a platform for common analyses to be performed.

## Results and Discussion

### Gathering information to develop a complete database centered on P. tricornutum genes

The DiatOmicBase website offers a gene-centered approach to facilitate exploration of the functional roles and regulation of diatom genes using omics-based resources. Considering *P. tricornutum* as a primary example, 12,392 gene pages corresponding to the latest annotation of protein coding genes (12,357 from the nuclear genome, 35 from the chloroplast, 60 genes from the mitochondria) contain sequence information gathered on a genome browser (Figure 1), alongside general descriptions related to their functional annotation and evolutionary history (Figure 2).

**Figure 1.**
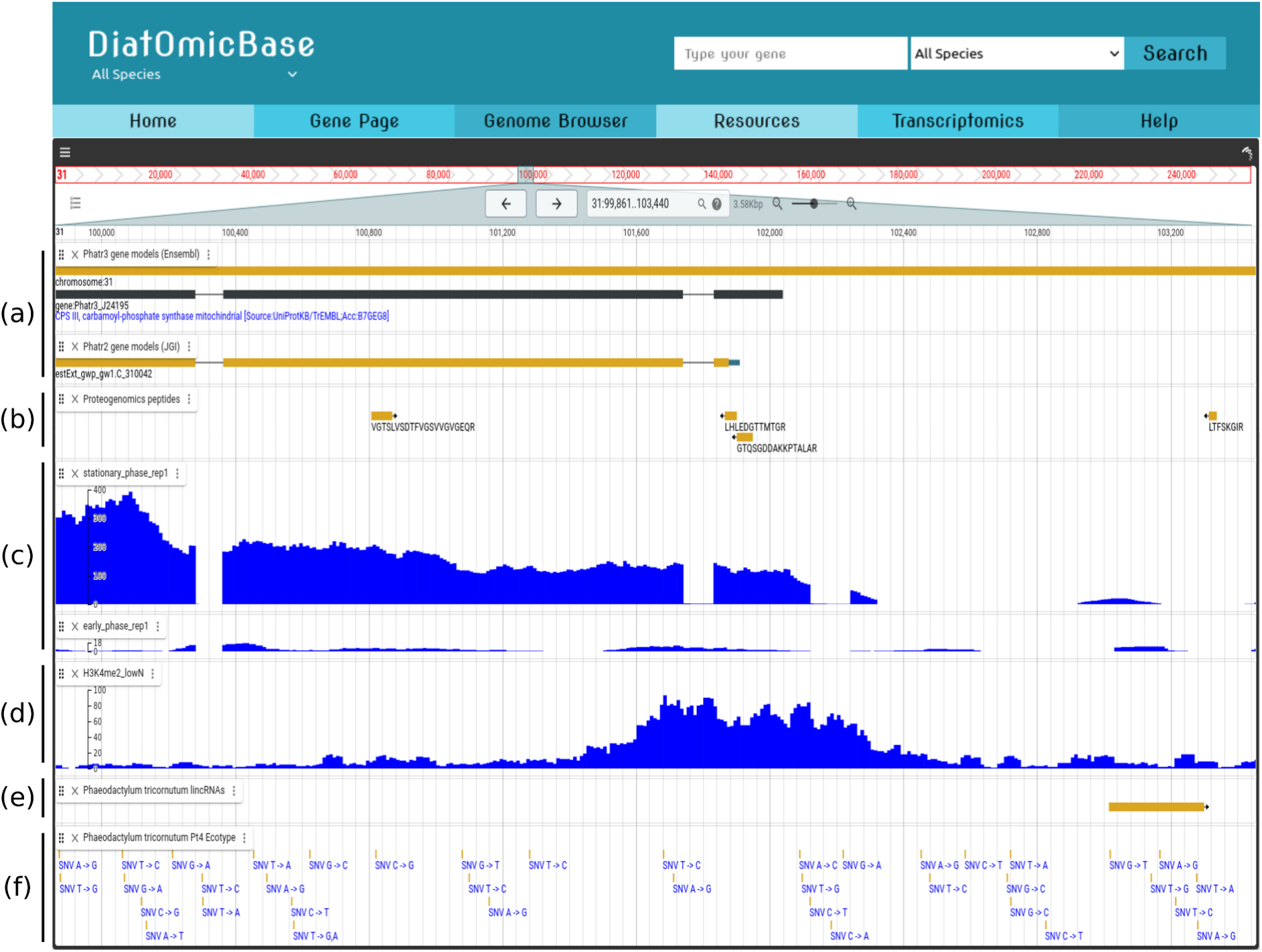
Snapshot of the DiatOmicBase genome browser illustrating the different features used for an integrative analysis of the CPS III gene (Phatr3_J24195). (a) Phatr3 and Phatr2 gene models can be compared at different zoom levels. (b) Zooming in on the 5’ region reveals two peptide sequences mapped using a proteogenomic pipeline on an exon predicted by the Phatr3 gene model but not by Phatr2. (c) Visualization of read mapping densities (here from Kwon et al., 2021) also validates the Phatr3 prediction. (d) Density plots of histone marks (H3K4 methylation) under nitrate-depleted conditions. (e) Long non-coding RNAs (Cruz de Carvalho et al., 2016) (f) Single nucleotide variants from 10 different ecotypes (Rastogi et al., 2020). This figure specifically shows only the Pt4 variant.

**Figure 2.**
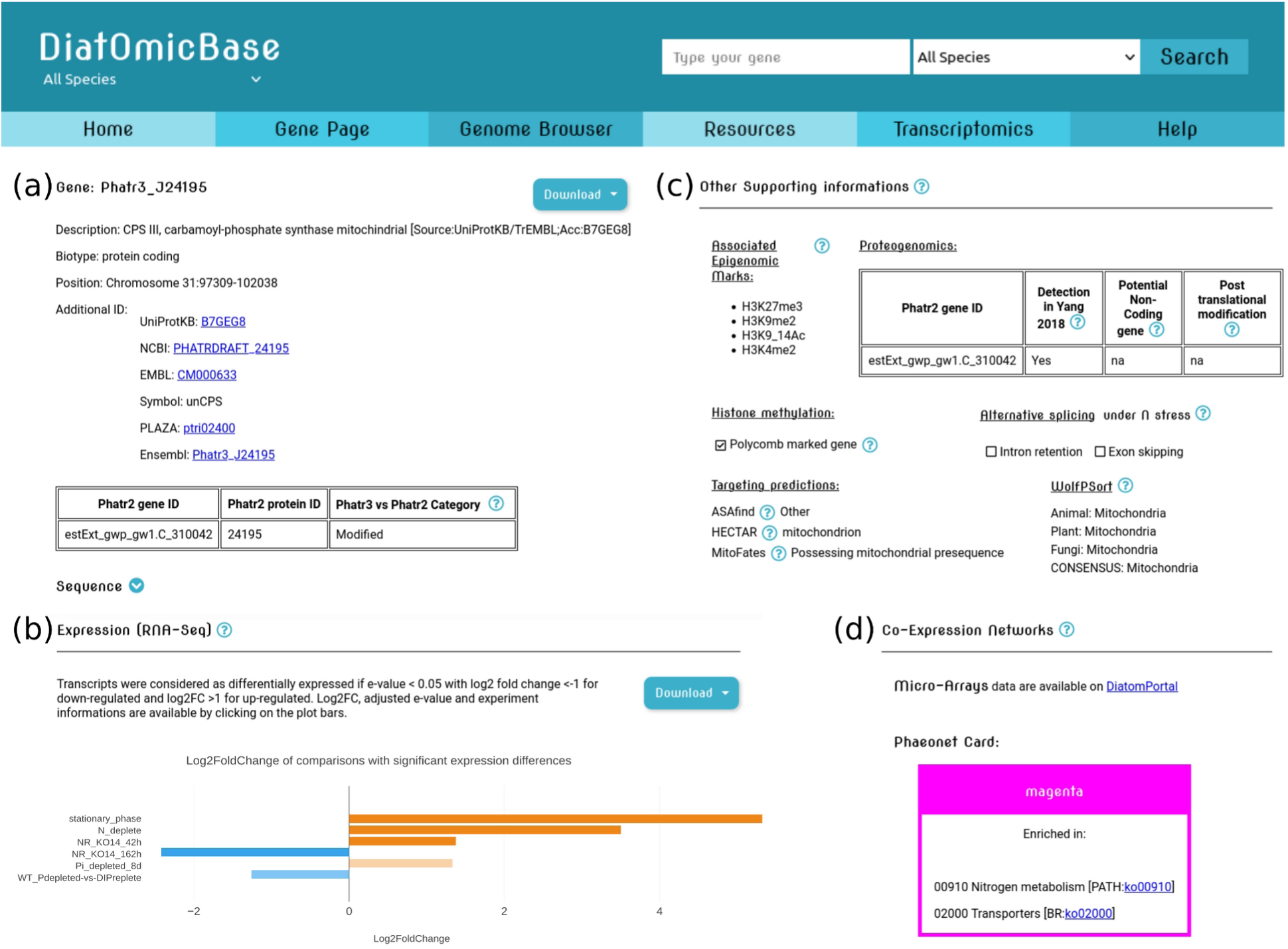
Snapshot of DiatOmicBase gene page for CPS III. (a) The different gene identifiers and links are provided to facilitate navigation between databases. (b) A barplot shows the log2 fold change of several RNA-Seq comparisons. The "stationary_phase" comparison contrasts early and stationary growth phases (Kwon et al., 2021). "N_deplete" compares nitrogen-free and nitrogen-replete samples after 48 [1] hours of treatment (Levitan et al., 2015). NR_KO14 refers to a transgenic knockout line for the nitrate reductase NR gene compared to wild-type cultures. In this experiment cells were grown in N free media and subsequently resuspended in NO3- for 42 and 162 hours (McCarthy et al., 2017). “Pi_depleted_8d” compares phosphate (Pi) replete cultures with cultures after 8 days of Pi depletion (Cruz de Carvalho et al., 2016). “WT_Pdepleted-vs-DIPreplete” compares wild-type strains grown in Pi depleted medium and replete with Dissolved Inorganic Phosphorus (DIP (Li et al., 2022). (c) The "Other supporting information" paragraph summarizes information collected from different studies (explanations and references are provided using the interactive "question mark"). Notably, HECTAR and Mitofates predict the expression of CPS III in the mitochondrion. (d) To further integrate functional data, a link is provided to access the corresponding microarray-based co-expression network and the Phaeonet clusters (based on RNA-Seq data). A snapshot of the complete page is available as Figure S2.

DiatOmicBase uses the 27.4 Mb **assembly** obtained in 2008 (Bowler et al., 2008) from the sequencing of *P. tricornutum* accession Pt1 8.6 (deposited as CCMP2561). As the latest published **gene prediction**, **Phatr3** (Rastogi et al., 2018) is defined as the standard in the website. This reannotation was obtained using existing gene models, expression data and protein sequences from related species to train prediction programs and predicts 12,089 gene models. To preserve the continuity of research by conserving gene identifier correspondence, the previous gene annotation, **Phatr2** (Bowler et al., 2008), comprising 10,402 gene models, is also available considering that several genes have been manually annotated on this version by the diatom research community. Only 4,667 Phatr3 gene models display a perfect correspondence with Phatr2. The other Phatr3 gene models can be new (1,489), have a modified 5′ and/or 3′ (4,709), can be merged (194), split (262), antisense (346), or required manual curation (566).

The genome browser also displays the annotation based on **proteomics** data. Yang et al., (2018) analyzed the proteome of 45 samples from *P. tricornutum* grown under eight different conditions. The peptide sequences resulting from the protein digestion and mass spectrometry analysis are shown on the genome browser. Using a proteomic pipeline integrating a dedicated protein search database, Yang et al., (2018) confirmed 8,300 Phatr2 genes, and identified 606 novel proteins, 506 revised genes, and 94 splice variants that can all be found on the genome browser. Discrepancies in gene model prediction between Phatr2 and Phatr3 can be resolved either using these proteomic data, or by analyzing mRNA expression and possible splicing variants using the multiple transcriptomics datasets centralized in the DiatOmicBase genome browser (see case study below, Figure 1 and Figure S1). To allow visualization of within-species gene diversity beyond the Pt1 8.6 strain, the database also includes genomic data from ten *P. tricornutum* ecotypes, covering broad geospatial and temporal scales (Martino et al., 2007), that have been re-sequenced (Rastogi et al., 2020). Variant calling results including SNPs, small insertions and deletions, can be visualized on the genome browser.

In addition to the sequence-related features displayed on the genome browser (Figure 1), several other descriptions are available on each gene page (Figure 2 and Figure S2). Gene pages are accessible by querying the database using Phatr2 or Phatr3 identifiers, terms or identifiers for GO, KEGG and Interpro databases. Keyword searches are available on the general search bar and on the gene page search. Finally, gene pages can be retrieved using nucleotide, protein and translated nucleotide (blastx) BLAST, with the possibility to tune various parameters.

The **gene annotations** include identifier correspondences with UniprotKB and NCBI, retrieved from their respective databases, and the former gene annotation Phatr2 provided in Rastogi et al., (2018). Annotations from the GO project (Ashburner et al., 2000) were retrieved from UniprotKB (The UniProt Consortium, release 2021_01). GO terms for functional analyses were generated via automatic annotation, and cover three domains: the Subcellular Component where the gene products are localized, the Molecular Function informing the main activities of the gene product, and the Biological Process, the set of molecular events involving the gene product. Several domain and protein family predictions can be retrieved from UniprotKB, integrating 20 different databases (e.g., InterPro, Pfam, Gene3D, TIGRFAMs, CDD; the complete list is available here). These databases integrate different automated and/or manually curated protein signatures to ease the identification of protein functions. Functional orthologs, as previously reported in Aït-Mohamed et al., (2020) using the KEGG orthology database (Kanehisa, 2002), provide insights about molecular functions using hierarchically-structured biochemical pathways.

**Transposable Elements** (TEs) represent around 75% of the detected repetitive elements in the Phatr3 genome annotation (Rastogi et al., 2018; Giguere et al., 2021). With their ability to insert into genes or regulatory sequences, TEs act as key players in the organisation and expression of the genome, contributing to phenotypic diversity and, ultimately to the species evolution (Abbriano et al., 2023). A specific track has been created on the DiatOmicBase genome browser to visualize their positions by transforming the flat tables produced in Rastogi et al., (2018) into gff files. The majority (2,790, ∼75%) of the TEs are associated with epigenetic marks (see below), which can also be important regulators of gene expression (Figure S2).

**Epigenetic modifications** play a pivotal role in regulating gene expression, influencing development, adaptation to environmental changes, and maintaining genome stability. In *P. tricornutum*, whole genome methylation (Veluchamy et al., 2013) and the distribution of five histone marks (Veluchamy et al., 2015) have been described for the most used *P. tricornutum* strain, Pt1 8.6 (CCMP2561) grown in standard culture conditions. The methylome was obtained by digestion of three replicate DNA samples with the methyl-sensitive endonuclease McrBC followed by hybridization to a 2.1-million-probe McrBc-chip tiling array of the *P. tricornutum* genome (Veluchamy et al., 2013). The 3,950 methylated regions shown on the genome browser result from normalization on these three biological replicates. Five histone marks were chosen as they are known to be involved in transcriptional activation or repression: H3K4me2, H3K9me2, H3K9me3, H3K27me3 and H3AcK9/14. Two biological replicates of each histone modification were analyzed using ChIP-Seq, resulting in the discovery of 119,000 regions annotated with a set of chromatin states covering almost 40% of the genome. Following this first description of a diatom epigenomic landscape, changes have been examined in response to nitrate depletion, both by analyzing histone modifications (H3K4me2, H3K9/14Ac and H3K9me3 using Chip-seq) and DNA methylation (bisulfite deep sequencing) (Veluchamy et al., 2015). These marks are also displayed on the genome browser. DiatOmicBase also provides a direct link to the PhaeoEpiView epigenome browser which include epigenomic data (mostly DNA methylation and post-translational modifications of histones) on the newly assembled genome (Filloramo et al., 2021).

Different kinds of **small noncoding (sn)RNAs** (25 to 30 nt-long) have been described in P. tricornutum (Rogato et al., 2014) after sequencing of short RNA fragments isolated from cells grown under different conditions of light and nutrients. Their sequences have been mapped onto the genome browser. The majority of snRNAs map to repetitive and silenced TEs marked by DNA methylation and recent evidence indicates their role in the regulation of epigenetic processes (Grypioti et al., 2024). Other snRNAs target DNA-methylated protein-coding genes, or are derived from longer noncoding RNAs (tRNAs and U2 snRNA) or are of unknown origin. Long noncoding (lnc)RNA sequences have been shown to play a significant role in transcriptional mechanisms and post-translational modifications (Statello et al., 2021; Mattick, 2023). Although most of the well characterized lncRNAs stem from mammalian systems, recent work has shown that marine protists, including diatoms, all express lncRNAs (Debit et al, 2023). Among the different categories of lncRNAs, over 1,500 lincRNAs (intergenic lncRNAs) and ∼3,200 lncNATs (antisense lncRNAs), that have previously been predicted from transcriptome mapping to the P. tricornutum genome (Cruz de Carvalho et al., 2016; Cruz de Carvalho & Bowler, 2020), are likewise available on the genome browser, but are not integrated on the gene pages.

Links to a range of already existing web databases for **comparative genomics** are also provided. Information can be mined using PLAZA Diatoms (Osuna-Cruz et al., 2020) that includes structural and functional annotation of genome sequences derived from 26 different species, including 10 diatoms. Complementary annotations, homologous and orthologous gene families, synteny information, as well as a toolbox enabling a graphical exploration of orthologs and phylogenetic relationships are available on the corresponding gene pages of PLAZA. Alongside this, **evolutionary history annotations** based on a ranked BLAST top hit approach obtained from 75 combined libraries from different taxonomic groups across the prokaryotic and eukaryotic tree of life have been generated (Rastogi et al., 2018).

**Co-expression networks** gather genes with similar expression patterns across samples, suggesting that they could be related functionally, regulated in the same way, or belong to the same protein complex or pathway. In *P. tricornutum*, two different studies of co-expression networks have been published. The first one, published by Ashworth et al., (2016) explored the hierarchical clustering of 123 **microarray** datasets generated from studies of silica limitation, acclimation to high light, exposure to cadmium, acclimation to light and dark cycles, exposure to a panel of pollutants, darkness and re-illumination, and exposure to red, blue and green light. A link to the corresponding gene page on DiatomPortal, the web platform containing this data, is available. More recently, Ait-Mohamed et al., (2020) performed Weighted Gene Correlation Network Analysis (WGCNA) of 187 publicly available and normalised **RNA-Seq** datasets generated under varying nitrogen, iron and phosphate growth conditions (Cruz de Carvalho et al., 2016; McCarthy et al., 2017) to identify 28 merged modules of co-expressed genes. The gene cluster identifier (denoted **PhaeoNet card**) and its KEGG pathway enrichments are indicated for each gene within these modules in DiatOmicBase. PhaeoNet modules for each gene are detailed in the supplementary information file available on the resource page.

Finally, supplementary information regarding the functions of each protein are provided: **post-translational modifications** extracted from the work of Yang et al., (2018); alternative splicing from the work of Rastogi et al., (2018) and also by analyzing the RNA-Seq data centralized in DiatOmicBase; *in silico* **targeting predictions** regrouping multiple different predictive tools: HECTAR under default conditions (Gschloessl et al., 2008); ASAFind using Signal P v 3.0 (Gruber et al., 2015; Dyrlov Bendtsen et al., 2004); MitoFates with cutoff threshold 0.35 (Fukasawa et al., 2015); and WolfPSort with animal, plant and fungal reference models (Horton et al., 2007), as previously assembled in Aït Mohamed et al., (2020).

Users have the possibility to leave comments on each gene page, indicating a reference and one or more predefined labels to facilitate indexing (Figure S2). This tool is expected to help the community to centralize information that is not easily accessible or cannot be automatically retrieved, such as the availability of transgenic lines, or evidence for transcriptional regulation or alternative splicing. A peer-reviewed reference is mandatory and comments can be moderated by the DiatOmicBase committee. The comments can also be used to specify a correct gene model when different gene annotations (Phatr2 and Phatr3) are not consistent.

### Transcriptomic Data and Analysis

We collected raw data available at NCBI from the RNA-Seq short reads BioProjects published from *P. tricornutum* (different ecotypes and selected transgenic lines), exposed to different conditions. These kinds of studies cover nutrient limitation (nitrate, phosphate, iron, vitamin B12), responses to chemical exposure (decadenial, naphthenic acids, glufosinate-ammonium, L-methionine sulfoximine, rapamycin, and nocodazole), different CO_2_ and light levels, response to grazing stress or competition. Samples involving transgenic lines (nitrate reductase and aureochrome photoreceptor knock-outs, alternative oxidase and cryptochrome knock-downs, chitin synthase transgenic cell lines, *etc*.) were compared to the corresponding wild-type samples. Morphotype-related transcriptomes were all compared together as well as growth stage transcriptomes. When studies consisted of time series, sample sets were compared in control *versus* treatment pairs for each timepoint but not between timepoints. As of July 2024, DiatOmicBase includes 33 studies, encompassing 1,398 samples and 135 comparisons.

For each BioProject, the most relevant pairwise comparisons were chosen to assess gene expression regulation in the different sample sets using the R package DESeq2 (Love et al., 2014). Bar plots showing up- and down-regulation illustrate the results of the pre-computed comparisons on each gene page. Only significant comparisons are graphically shown but the list of non-significant comparisons and samples without read matches are provided.

On the transcriptomics page, users can also reanalyze public Bioprojects, defining the samples to compare (see case study 2). For public data, the gene-level read counts are already computed for each sample, and users only have to select the samples to compare. Moreover, users can also analyze their own data (see case study 3). For private data, inputs are a gene-level read count table or any equivalent expression matrix and a table informing how the samples should be grouped to be compared.

Gene expression analyses can subsequently be performed using the web application “integrated Differential Expression and Pathway analysis” (iDEP; Ge et al., 2018). Connecting several widely used R/Bioconductor packages and gene annotation databases, iDEP provides a user-friendly platform for comprehensive transcriptomics analysis, including quality control plots, normalization, PCA, differential expression analysis, heatmaps, pathway and GO analysis, KEGG pathway diagrams, functional annotation, co-expression networks, interactive visualizations, and downloadable gene lists. iDEP enables pairwise comparisons or more complex statistical models including up to 6 factors. Co-expression networks can be examined using WGCNA. All the analysis steps are customisable; methods and parameters can be easily tuned using dialog boxes.

Finally, expression patterns of several genes can be analyzed by drawing customized heatmaps (Figure 3). In this case the user can provide a gene list and select the sample comparisons to be shown on the plot.

**Figure 3.**
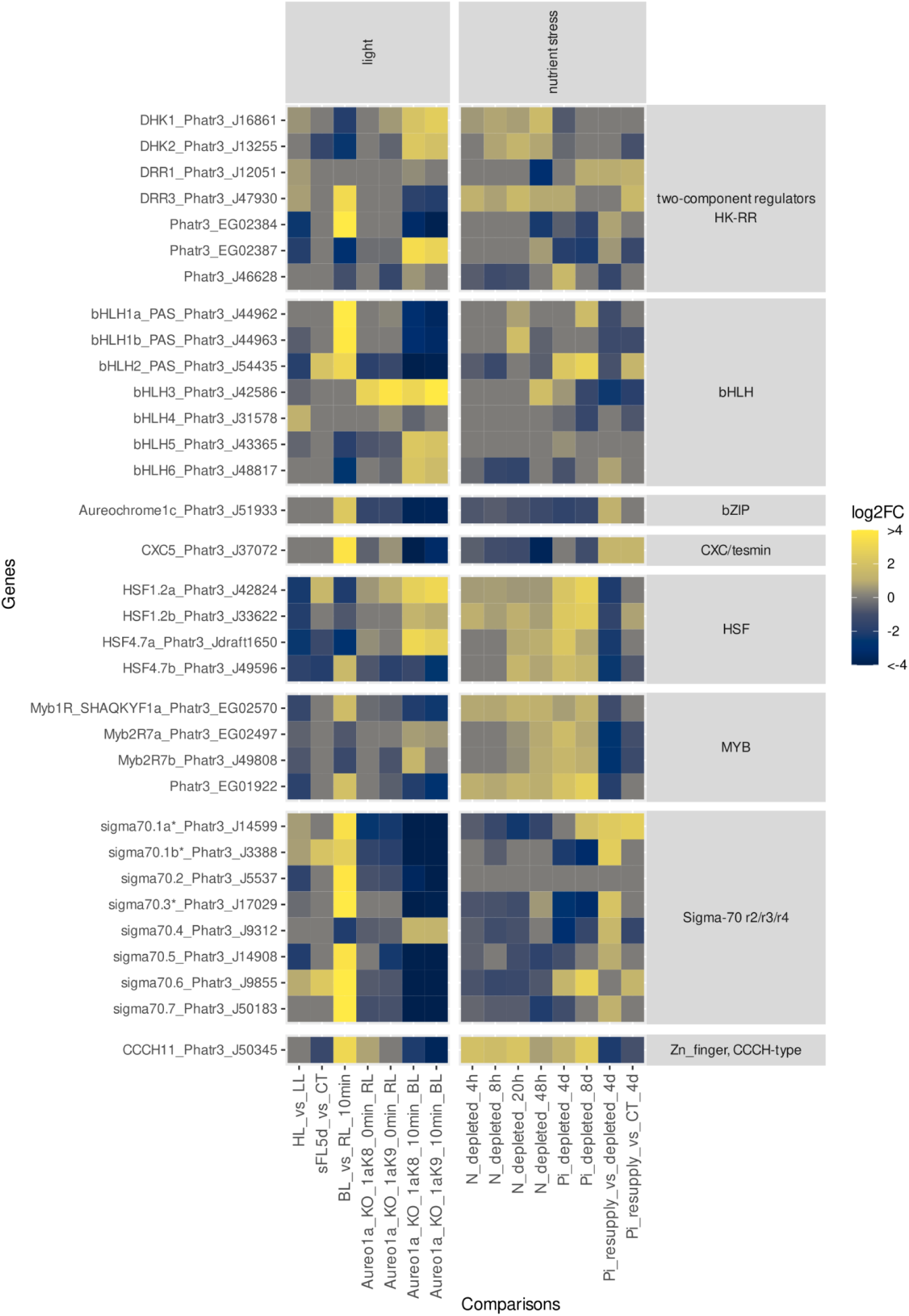
RNA-Seq heatmaps of two components regulators and transcription factors Heatmap showing gene expression differences in light and nutrient stress experiments for different genes of interest. HL_vs_LL compares low light and high light acclimated wild-type (WT) strains (Agarwal et al., 2022). sFL5d_vs_CT compares strains acclimated to constant light conditions to severe light fluctuation conditions (Zhou et al., 2022). BL_vs_RL_10min compares WT strains under red light (RL) and 10 minutes after a shift to blue light (BL); In samples KO_1aK8_10min_BL and KO_1aK9_10min_BL, Aureo1a knockout K8 and K9 strains are compared to WT under RL, and after a 10 minutes shift to BL (Mann, et al., 2020). N_deplete_ compares nitrate-deplete and control samples 4h, 8h and 20h after transfer to nitrogen-replete medium (Matthijs et al., 2016). N_deplete_48h compares nitrogen-free and nitrogen-replete samples after 48h of treatment (Levitan et al., 2015). Pi_depleted_4d and Pi_depleted_8d compare control samples and cultures without phosphate supplement during 4 and 8 days. Pi_resupply-vs-depleted_4d and Pi_resupply-vs-CT_4d compare samples with phosphate re-supplementation after 4 days of starvation (Cruz de Carvalho et al., 2016). * based on the prediction supported by the database, these genes are localized in the plastid.

### Case studies

#### Improving gene model prediction and visualization of gene expression and regulation

DiatOmicBase can aid functional analyses of diatom genes by improving prediction of protein-coding genes and their regulation. In Figures 1 and 2, we show the information that can be readily obtained in DiatOmicBase, using as an example the *P. tricornutum CPS III* gene, encoding a carbamoyl phosphate synthase involved in the urea cycle. In diatoms, the discovery of an ornithine-urea cycle (OUC) was unexpected as it was previously thought to be specific to animals, to allow removal of excess NH4^+^ derived from a protein-rich diet.

Rather than eliminate NH_4_, diatoms have been proposed to conserve this precious resource by using the pathway to cope with fluctuations in nitrogen availability in the ocean (Allen et al., 2011, Smith et al., 2019), maintaining the cellular balance of carbon and nitrogen. The key enzyme of the OUC, carbamoyl phosphate synthase (CPS) is encoded in the *P. tricornutum* genome by two copies. One copy, named *unCPS* or *CPS III* (Phatr3_J24195) and using ammonium as a substrate, is predicted both by HECTAR and MitoFates to be targeted to the mitochondria (Figure 2c), in agreement with the mitochondrial localization observed by microscopy (Allen et al., 2011). Furthermore, a *T. pseudonana* homologue of *unCPS* (Thaps3a_40323) has been recovered from proteomic data derived from purified mitochondrial fractions. A second *P. tricornutum* paralog, pgCPS2 (encoded by Phatr3_EG01947) has been inferred to be cytosolic.

In DiatOmicBase, the peptide alignment from proteogenomic studies confirmed the Phatr3 gene model of *CPS III*, compared to the Phatr2 gene model (estExt_gwp_gw1.C_310042; Figures 1, 2 and S1), predicting a protein extended by 133 amino acids at the N-terminus, due to the translation from an earlier ATG initiation site in the genome (chromosome 31 position 102308 reverse strand, c.f. chromosome 31 position 101909 reverse strand). This gene model and also intron position were confirmed by analyzing RNA-Seq data from conditions where the *CPS III* gene is strongly expressed (Figure S1).

Considering quantitative gene expression trends, reanalysis of transcriptomic data in DiatOmicBase has shown that *CPS III* is highly expressed in cells experiencing three days of nitrogen deprivation (Levitan et al., 2015). Interestingly, its expression is also induced under phosphorus (P) depletion conditions, whereas Pi repletion of starved cells reverse this trend. These responses to Pi availability could be the result of the rapid cross-talk between Pi and N metabolism observed in diatoms (Helliwell et al., 2021). *CPS III* was also overexpressed in a nitrate reductase (NR) knockout line compared with the wild-type (wt) under conditions of nitrogen repletion (e.g., comparing the response of NR knockout versus wt in cells repleted with nitrogen for 42 h), but was down-regulated when cells became limited for this nutrient (e.g., NR KO versus wt after 162 h of nitrogen resupply) (McCarthy et al., 2017; Figure 2b). *CPS III* is also down-regulated in cell lines incubated in the complete absence of nitrogen compared with nitrogen-replete media for 4 and 20 hours (Matthijs et al., 2016), but was up-regulated when cells experienced prolonged nitrogen starvation (Levitan et al., 2015). It is therefore possible that nitrate uptake and internal cellular nitrate concentrations play a role in the hierarchical regulation of *CPS III* expression. We also note a dramatic increase in *CPS III* expression in stationary-phase cells compared with exponential-phase cells, which may relate to functions in amino acid scavenging and recycling, but also in nutrient starvation responses. Information on *CPS III* gene expression in additional growth conditions and physiological states can be obtained by analyzing other centralized omics data (e.g., Figure S2).

The histone marks H3K4me2 and H3K9/14Ac, consistent with transcriptional activity, mapped to the 5’ region of the gene both in nitrate replete and deplete conditions, suggesting that CPS III is also important in nitrate replete cells (Figure 2c). We furthermore note that *CPS III* groups in the PhaeoNet magenta card, a module of co-expressed genes that appear to be enriched in functions implicated in organelle amino acid and nitrate metabolism (Figure 2d). Other members of this module include the plastidial glutamate synthetase (Phatr3_J50912) and N-acetyl-gamma-glutamyl-phosphate reductase implicated in the plastidial ornithine cycle (Phatr3_J36913), as well as several other genes encoding proteins involved in nitrate assimilation. This might be consistent with a role of the urea cycle under nitrate replete conditions in recycling excess assimilated plastidial amines. Finally, while no TEs were found in the coding region, a lincRNA was detected in the 5’ region of the gene. A dozen small RNAs were mapped onto the gene, and ecotypes displayed between 7 and 31 single nucleotide variants (Figure S2), suggesting further possible hierarchical factors influencing the function and evolution of this gene.

#### Identification of common or distinct regulators of diatom responses to light and nutrient variations

Here we show how transcriptomic resources centralized in DiatOmicBase can be exploited to perform novel comparative functional analyses across multiple datasets (i.e., meta-analyses) and derive new information about common or specific regulators possibly implicated in diatom acclimation to environmental cues. Fluctuations in nutrient availability are recurrent in the marine environment, with nitrogen (N) and phosphorus (P) being amongst the main limiting nutrients for primary productivity. Both nitrogen and phosphorus deficiencies lead to a reduction in photosynthetic activity and a halt in cell division in diatoms (Jaubert et al., 2022; Levitan et al., 2015; Cruz de Carvalho et al., 2016; Matthjis et al., 2016). This may result in a cellular quiescent state, which is reversed when nutrients are resupplied (Cruz de Carvalho et al., 2016; Matthjis et al., 2016; Dell’Aquila and Maier, 2020). On the other hand, light is the major source of energy for photosynthesis, and a critical source of information from the external environment. In recent years, the exploration of diatom genomes and of transcriptomic data obtained from cells exposed to different light conditions (wavelength, intensity and photoperiod cycles) and targeted functional analyses of selected genes in *P. tricornutum* has identified peculiar regulators of diatom photosynthesis (Lepetit et al., 2022), novel photoreceptors responsible for light perception, and an endogenous regulator controlling responses to periodic light-dark cycles (Jaubert et al., 2022). However, the regulatory cross-talk between light and nutrient signaling pathways remains poorly understood in diatoms. To this aim, here we reanalyzed already available RNA-Seq data from cells experiencing different light conditions (high light stress, fluctuating light, different colours) as well as different nitrate and phosphate depletion and repletion conditions.

In Figures 3, S3 and S4, we focused on gene expression changes for two-component regulators and transcription factors (TFs) encoded in the *P. tricornutum* genome. These regulators orchestrate the physiological response(s) to a given stress by participating in signaling cascades through differential expression regulation. The genome of *P. tricornutum* contains a high number of bacterial-like two-component sensory histidine kinases (Bowler et al., 2008), defined on the basis of the presence of a histidine kinase domain (PF00512) and/or a response regulator domain (PF00072). In addition, the histidine phosphotransmitter protein (Hpt), which was reported missing in the first version of the genome (Bowler et al., 2008), was later identified in Phatr3 (Phatr3_J33969, PF01627) and was included in the analysis (Figure 3). Regarding two-component regulators, we observed significant gene expression changes for some of these regulators following a red to blue light shift (e.g., *DHK1, DHK2, DRR3, EGO2384, EGO2387*). The loss of these responses in two independent Aureochrome blue light receptor KO lines (Mann et al., 2020) compared to wild-type cells indicates that this blue light photoreceptor participates in the regulation of their expression. By contrast, *Phatr3_J46628* is not affected by any changes in light conditions, but is over-expressed under prolonged phosphate depletion (4d) (Figure 3). Interestingly, *DDR3* is found to be up-regulated under blue-light treatment, but also under both N (from 4h to 48h) and Pi depletion (4 days) as well as under Pi resupply (Figure 3 and S3), but not under HL. It therefore appears likely that a blue light-activated signaling cascade participates via DDR3 in the regulation of cellular responses to nutrient availability, but not in photoprotection. *Phatr3_EG02384* is expressed in the opposite fashion in response to HL and nitrogen depletion, suggesting a possible antagonistic role in light and nutrient signaling pathways. On the other hand, *DRR1* appears not to be affected by light signals, but is specifically expressed under prolonged N deficiency and Pi depletion.

The analysis of TF expression patterns across treatments also provides interesting information (Figures 3 and S4). Between the genes encoding bHLH domain-containing proteins, the bHLH1a-PAS also known as RITMO1 shows a rapid induction by blue light treatment, which is repressed in *Aureo1* photoreceptor mutants, as previously shown (Mann et al., 2020). Neither HL treatments, nor short N and Pi deficiency affect its expression, as expected considering the role of this protein as an endogenous and robust timekeeper (Annunziata et al., 2019). Its expression and possible activity is, however, affected by prolonged nutrient stress treatments. bHLH2-PAS is differentially expressed under the tested light conditions as well as Pi deficiency, while bHLH3 appears to be expressed specifically under N and Pi deficiency. Interestingly, its expression is not induced by blue light, but is significantly affected in the *Aureo1a* mutants under red light. It is thus possible that *Aureo1a*, which is also a TF with a bZIP domain, can participate in regulation to nutrient changes, independently of its activity as a blue light sensor. A light-independent activity of *Aureo1a* is also supported by a recent study reporting altered rhythmic gene expression in *Aureo1* mutants compared to wild-type cells in constant darkness (Madhuri et al., 2024). Of the genes encoding sigma 70 factors, it is interesting to note that their expression is strongly modulated by HL or blue light, and for the three factors predicted to be plastid localized (sigma 70.1a, 1.b and 70.3*) also by Pi availability. This is consistent with a possible role for these factors in regulating plastid gene expression, which could also be sensitive to the physiological state of this organelle. Other TFs modulated by nutrient availability and blue light include Aureochrome1C of the bZIP family and Pt-CXC5 of the CXC/tesmin family, both of which are downregulated under nutrient stress and upregulated upon nutrient resupply, namely P, as well as under blue light (Figure 3). On the other hand, the opposite trend can be found for TFs of the HSF (HSF1.2a/b; HSF4.7a/b) and Myb families, which are also modulated by nutrient status and light, being up-regulated under nutrient and high light stress (Myb2R7a/b), as well as blue light (Myb1R_SHAQKYF1a; Phatr3_EG01922).

This analysis reveals the complex cross-talk between the regulatory pathways operating under light and nutrient fluctuations, which could enable opening new exploratory and functional research avenues aiming to shed light into the molecular processes underpinning diatom resilience to environmental stresses.

#### DiatomicBase for the *de novo* analysis of previously unpublished RNA-Seq data

In DiatomicBase, we have also developed the possibility for users to analyze their own RNA-Seq data. Here, we show the analyses of new data obtained from *P. tricornutum* lines over a progressive two-week iron (Fe) starvation time-course. More specifically, we compared gene expression changes between cell lines grown in Fe replete media and those experiencing short-, medium- and long-term Fe withdrawal conditions (Figure S5).

Principal Component Analysis of the RNA-Seq data, performed with the integrated iDEP platform in DiatOmicBase, revealed three groups, corresponding to Fe-replete, short-term (3 days), and medium/ long-term (7, 14 days) Fe limitation (Figure 4a). Calculation of the k-means revealed four distinct clusters enriched in different GO terms that were differentially regulated in response to different periods of Fe limitation (Figure 4b). The first cluster (generated as Cluster A by the iDEP calculations) was strongly induced after 3 days Fe withdrawal relative to Fe-replete conditions and was significantly (P< 10^-15^) enriched in genes encoding proteins with functions relating to photosynthesis (photosynthesis light reactions, protein-chromophore linkage, generation of precursor metabolites and energy), alongside translation and peptide biosynthesis-associated processes (Figure 4c). This might relate to a transcriptional and translational upregulation of light-harvesting protein complexes in particular to compensate for diminished Fe availability limiting the synthesis of photosystem I (Gao et al., 2021). This contrasted with a second cluster (Cluster C) that was uniquely downregulated in short-term Fe limitation conditions, and showed a weaker (P < 0.001) enrichment in photosynthesis and transcriptional regulation processes but no other enriched GO terms (Figure 4b). The final two clusters, Clusters B and D, were strongly up- and downregulated, respectively, by medium and long-term Fe limitation compared to short-term and Fe-replete conditions. These showed weak enrichments (P < 0.0001) in transport processes, i.e., ion and organic metabolite transport (Figure 4b). These may relate to the induction of Fe uptake systems in response to prolonged Fe withdrawal, alongside changes in the internal metabolite profiles and organic acid transport activities of Fe-limited cell lines (Gao et al., 2021).

**Figure 4.**
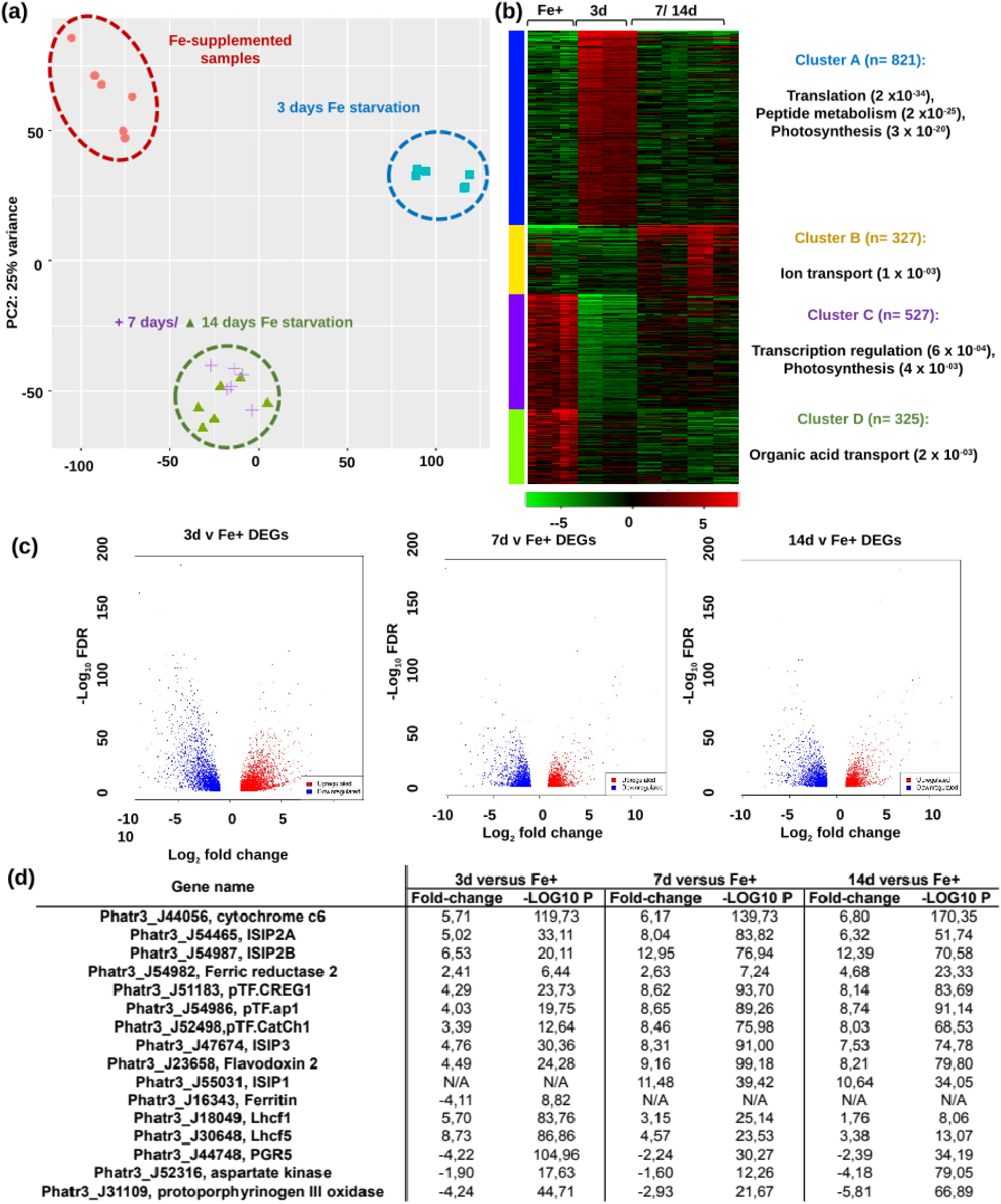
de novo RNA-Seq analysis of iron starvation response. Comparison of gene expression from cells adapted to 12h light:12h night cycles under 19°C, and either Fe-replete (“Fe+”), 3 day (“FeS”), 7 day (“FeM”) or 14 day (“FeL”) Fe-limitation. A schematic experimental design is provided in Figure S5. (a) PCA of RNA-Seq Transcripts Per Million (TPM) values calculated with the integrated iDEP module in DiatOmicBase. Three distinct treatment groups are visible, suggesting relative equivalence of FeM and FeL treatments. (b) k-means clusters, showing selected enriched GO terms (P < 0.001), calculated with the iDEP module in DiatOmicBase. Four clusters with distinctive biological identities show different relationships to both short-term and medium/ long-term Fe-limitation. (c) Volcano plots of DEGs inferred with iDEP with threshold P-value 0.1 and fold-change 2. (d) Tabulated fold-changes and −log10 P-values of genes associated with Fe stress metabolism, following Gao et al., (2021).

While the data from DiatOmicBase provide insights into transcriptomic changes, they can be used to inform user construction of more complex evaluations of the links between gene expression and function. As an example of this, in Figure 4c we present Volcano Plots for DEGs generated from the DiatomicBase site, alongside tabulated fold-change and P-values calculated for genes known to be involved in Fe stress metabolism (Figure 4d; Gao et al, 2021). Concerning known Fe-stress associated proteins, many were already strongly upregulated after three days Fe limitation compared to Fe-replete lines, confirming the immediate impacts of Fe withdrawal on cell physiology (Figure 4c) (Gao et al., 2021). The most significantly upregulated of these genes at all three time points (P-value < 10^-100^ for 3, 7 and 14-day Fe limitation) encoded a plastidial cytochrome c_6_ (Phatr3_J44056), which mediates photosynthetic electron transfer between cytochrome b_6_f and photosystem I, underlining the importance of photosystem remodelling in both short-term and sustained Fe limitation (Allen, de Paula et al., 2011). We note that *P. tricornutum* does not possess an endogenous plastocyanin gene (Groussman, Parker et al., 2015) that could be induced to compensate for cytochrome c_6_ under Fe-limitation.

Consistent with previous studies, we see the upregulation of known iron-stress related genes at all Fe-limitation time points studied. Constitutively upregulated genes include *ISIP2a/ phytotransferrin* (*Phatr3_54465/ Phatr3_54987*) and *ferric reductase 2* (*Phatr3_J54982*), implicated in the reductive uptake of Fe from the cell surface (Morrissey, Sutak et al., 2015, McQuaid, Kustka et al., 2018); alongside genes encoding the newly identified plastid-targeted partners of ISIP2a *pTF.CREG1* (*Phatr3_J51183*), *pTF.ap1* (*Phatr3_J54986*), and the gene encoding the plasma membrane-located ISIP2a-binding protein pTF.CatCh1 (*Phatr3_J52498*) (Allen, Laroche et al., 2008, Turnšek, Brunson et al., 2021). These results suggest that reductive Fe uptake in diatom cells is relevant to both short- and long-term Fe limitation. We also observed upregulation at all time points for *ISIP3* (*Phatr3_J47674*), encoding a protein of unknown function but proposed to participate in the intracellular trafficking or storage of iron (Behnke and LaRoche 2020, Kazamia, Mach et al., 2022); and flavodoxin 2 (*Phatr3_J23658*), which can diminish cellular iron requirements of flavodoxin in photosystem I (Setif 2001, Lodeyro, Ceccoli et al., 2012).

Our data also provide insight into Fe-stress associated diatom genes that show more distinctive responses to Fe limitation. For example, ISIP1 (*Phatr3_J55031*), implicated in the non-reductive transport of Fe-siderophore complexes from the cell surface to the chloroplast, shows no evidence of induction in the three day treatment, but strong induction in medium- and long-term limitation datasets (Figure 4c) (Kazamia et al., 2018), suggesting that it is induced more slowly than reductive iron uptake strategies. This might reflect the greater energetic cost or gene coordination required by diatoms to produce siderophores in response to Fe limitation, whose synthesis pathway remains unknown. Most dramatically, the gene encoding the proposed plastid iron storage protein ferritin (*Phatr3_J16343*) showed no response to either medium or long-term Fe-limitation but was significantly downregulated (P < 10^-08^) in response to short-term Fe withdrawal. The role of ferritin in diatoms has historically been unclear, with some studies suggesting that it facilitates long-term Fe storage and tolerance of chronic starvation (Marchetti et al., 2009); and others that it may be transiently upregulated in response to Fe enrichment, allowing competitive removal of iron from the environment (Lampe et al., 2018, Cohen et al., 2018). Our data are broadly more consistent with the latter role for the *P. tricornutum* ferritin, although we note it is phylogenetically distinct to other diatom ferritins and may confer different physiological roles (Gao et al 2021). A greater similarity of this *P. tricornutum ferritin* gene with sequences from *Nannochloropsis* and ciliates than from other diatoms is also observed by analyzing PLAZA Diatoms.

Finally, our data provide some insights into the broader physiological responses that mediate short- and long-term diatom responses to Fe deprivation. The importance of light-harvesting complexes for short-term responses to Fe-limitation (Figure 4c) is underlined by the strong induction of genes encoding two Lhc proteins (Lhcf1, *Phatr3_18049*; Lhcf5, *Phatr3_J30648*) that directly interact with one another in the PSI LHC (Joshi-Deo et al., 2010), although are also both found in PSII (Gundermann et al., 2013; Nagao et al., 2021), and are implicated in low-light adaptation (Gundermann et al., 2013). In contrast, we observed strong downregulation of PGR5 (*Phatr3_J44748*), debatably implicated in cyclic electron flow around PSI in diatoms (Grouneva et al., 2011, Johnson et al., 2014). It has recently been proposed that *Chlamydomonas reinhardtii* and plant PGR5 indirectly participate in the delivery of Fe to PSI, which might explain its immediate sensitivity to Fe depletion in our data (Leister et al., 2022). In contrast, under long-term Fe-limitation but not under short- or medium-term conditions, we identified dramatic (P < 10^-50^) downregulation of the gene encoding Aspartokinase (*Phatr3_J52316*), involved in lysine biosynthesis, as well as the gene encoding Protoporphyrinogen oxidase (*Phatr3_J31109*), involved in the tetrapyrrole branch of chlorophyll/haem biosynthesis, which may point to an overall quiescence of core organelle metabolic pathways in response to sustained Fe deprivation (Allen et al., 2008; Ait-Mohamed, Novák Vanclová et al., 2020).

### Perspectives

DiatOmicBase provides a comprehensive database and analytic modules integrating genomic, epigenomic, transcriptomic and proteomic data from published diatom genomes. In this work, we focused on the resources centered around *P. tricornutum*. The first objective of this centralized database is to provide the community with a powerful tool for assessing correct gene models. Automated gene annotation remains error-prone, especially considering the large proportion of diatom genes whose function is unknown and which have no clear homologues in other systems. The peptide and RNA/DNA sequence mapping provided in DiatOmicBase may help to identify the correct form, as shown for the *CPS III* gene; DiatOmicBase makes provisions for user annotation and correction of individual gene pages. Given that *P. tricornutum* is the diatom species with the largest collection of genomic resources and data, the correct gene annotation in this species also represents a powerful support for correct gene prediction in other diatom species and from environmental data.

Analyzing the expression of genes in different circumstances has been facilitated by grouping results of several RNA-Seq experiments involving different conditions. In the case studies described in this work, we have shown how our tools can help to identify common or specific regulators of responses to various light changes and nutrient stress conditions. This offers new opportunities to characterize the still largely unknown signalling pathways involved in the perception and acclimation to complex changing environments. Using the iDEP pipeline, public RNA-Seq datasets can be re-analyzed, enabling to answer questions different from the ones in the original articles. In order to enrich our resources, we would like to encourage the community to inform us when new data becomes available.

DiatOmicBase also facilitates the study of gene functions, essential for deciphering the physiology and metabolic capacities of these microalgae and harnessing their potential for biotechnology and synthetic biology Assigning a phenotype to a protein requires overcoming many challenges (Heydarizadeh et al., 2014) as automated annotations should be validated by biochemical evidence and comparative analyses between wild-type and mutant lines. Metabolic engineering and synthetic biology resources in diatoms are rapidly evolving (Russo et al., 2023), yet they are limited by some aspects of diatom metabolism that are not yet well understood. For example, we still do not know how many metabolic pathways, including those that are important for biotechnology such as the biosynthesis of carotenoids and terpenoids, react to environmental conditions. Nor do we know the mechanisms by which these pathways are regulated, or the subcellular location of the different enzymes. Protein targeting predictions, expression data available or newly generated and analyzed in DiatOmicBase, in conjunction with metabolomics data can be exploited for strain engineering strategies, by improving gene-enzyme-function associations, identifying unknown enzymes, and reveal still elusive aspects of pathway regulation. For designing pathway engineering and optimizing cultivation strategies, DiatOmicBase can be particularly useful for investigating co-regulation of industrially-relevant metabolic pathways or key pathway nodes with transcription factors (Figure 3) across various conditions to identify regulators that can control entire metabolic pathways. These are primary genetic targets for strain engineering strategies which are promising and feasible (Song et al., 2023), but also remains a largely untapped strategy in diatom biotechnology.

The future development of the DiatOmicBase should include improved genome assemblies and annotations. Unfortunately, *P. tricornutum* genome complexity (especially TEs) has prevented significant improvement of the reference genome obtained by Sanger sequencing. A new assembly using the latest technologies of long read sequencing has recently been published (Filloramo et al., 2021). However, this long-read derived assembly still lacks continuity resulting in more than 200 scaffolds (while the Sanger assembly comprised 88 scaffolds with 33 chromosome-level scaffolds). As no gene annotation was associated with this assembly, it was not possible to fully exploit this new resource in this first version of DiatOmicBase, but we aim to include it following it’s annotation. As shown for the *CPS III* gene, DiatOmicBase should help to resolve conflicting gene model predictions derived by new sequencing and assembly projects. The database makes provisions for user annotation and correction of individual gene pages.

The addition of new diatom species in DiatOmicBase is also planned, especially those with the most developed omic resources. As of September 2024, preliminary DiatOmicBase portals are available for the model centric species *T. pseudonana* (https://www.diatomicsbase.bio.ens.psl.eu/genomeBrowser?species=Thalassiosira+pseudonana) and the model sexual diatom *P. multistriata* (https://www.diatomicsbase.bio.ens.psl.eu/genomeBrowser?species=Pseudo-nitzschia+multistriata). Users can automatically search for gene information in these different species. Further developments will include the addition of new proteomic data, the incorporation of comparative genomic and automated phylogenetic analyses of individual genes, functional genomic information from transgenic lines, and precomputed biogeographical distributions of environmental homologues of individual diatom genes from *Tara* Oceans (Vernette et al., 2022).

## Experimental procedures

### Website architecture

The back-end server of the website consists of an API server coded with the FastAPI framework. It includes a local PostgreSQL database that is accessed using the SQLAlchemy library. Data from NCBI/EBI/Ensembl are fetched and inserted or updated in the local database using a python loader script. The genome browser used is Jbrowse 2 (Buels et al., 2016). A R-shiny iDEP instance (Ge et al., 2018), hosted on the local server, was modified to contain *P. tricornutum* genome annotation and to automatically load public data from DiatOmicBase. A background worker coded in python is used to run more computationally intensive user calculations (Blast or differential expression analysis) and ensures that these do not use more resources than allowed, using a queuing system. The possibility to comment gene pages used Commento widgets. To ensure data privacy, Commento and its postgreSQL database are self-hosted.

The front-end part is developed with the React framework and uses static rendering with Next.js for performance. The FastAPI and Next.js instance communicates with the user through an Apache proxy to retrieve user requests and display results. Cite the Github address

#### Analysis of publicly available transcriptomic data

We collected raw data available at NCBI from the RNA-Seq short reads BioProjects published from *P. tricornutum* (different ecotypes and selected mutants), exposed to different conditions. Short read sequences were 35 to 150 bp long, principally generated from Illumina or DNBSeq. Fastq files were analyzed using the Bioinformatic pipeline nf-core/rnaseq (Patel et al., 2021). Briefly, raw reads were cleaned and merged before being mapped with Hisat2 on their reference genome and quantified using featureCounts (Liao et al., 2014), an algorithm specifically designed to quickly and efficiently quantify the expression of transcripts using RNA-Seq data. Read coverage files are displayed on the genome browser of each gene page.

For each BioProject, pairwise comparisons were chosen to assess gene expression regulation in the different sample sets using the R package DESeq2 (Love et al., 2014). All the replicates deemed to be available and of good quality (based on mean quality score computed with FastQC) were used to estimate biological and technical variation. In studies investigating transcriptomic responses to environmental variations, the tested conditions were compared with their respective control groups. In cases involving mutant lines, precomputed comparisons were made between mutants and wild-types.

Gene expression analyses can be performed with the web application “integrated Differential Expression and Pathway analysis” (iDEP, Ge et al., 2018). Data can first be explored using heatmap, k-means clustering, and PCA. Differential expression analysis can be conducted through two different methods; limma and DESeq2 packages and several visualization plots are available (e.g., Venn diagrams, Volcano plots, genome maps). Based on Ensembl annotation, pathway enrichment can be performed from GO and KEGG annotation using several methods: GSEA, PAGE, GAGE or ReactomePA.

#### RNA-Seq data generated in this study over a two-week iron (Fe) starvation time-course

Wild-type *P. tricornutum* v 1.86 cells were grown in ESAW medium at 19°C (Dorrell, PNAS 2021), under 12hLight:12Dark cycle (50uE light). Fe-replete (Fe+) and Fe-depleted (Fe-) ESAW media were both produced using iron-free reagents based on a protocol from Xia Gao (Gao et al., 2019). For the transcriptomic analyses, three different Fe limitation conditions were designed: three days Fe limitation as a short-term Fe-treatment (FeS), one week Fe limitation as a medium-term Fe-treatment (FeM) and two weeks Fe limitation as a long-term Fe-treatment (FeL). Fe-replete conditions (Fe+) were used as a control. RNA-Seq analysis were performed using two genetically distinct lines of *P. tricornutum* wild-type cells, transformed with pPhat and HA-Cas9 vectors without guide RNAs (Dorrell et al, 2024), to allow subsequent comparison of gene expression responses to mutants (data not shown). Reproductible transcriptome dynamics were observed for each culture, suggesting insertion of the pPhat and Cas9 vectors did not intrinsically bias Fe metabolism in these strains. Three biological replicates were performed for each cell line and condition. Cells were collected for the analyses at the exponential phase, and no other nutrient stress than Fe limitation influenced the results. Fast Fv/Fm (with 10% FP, 70% SP) tests were performed for all cell lines using a PAM (PAR-FluorPen FP 110, Photon Systems Instruments) prior to sampling. Fv/Fm values under Fe-replete values were measured at mean 0.62, suggesting that neither N nor Pi were growth-limiting in the media. Measured Fv/Fm values after 3 days Fe starvation were 0.55, suggesting Fe limitation of photosystem activity.

Total RNA was extracted from around 50 mg cell pellets by using the TRIzol reagent (T9424, Sigma-Aldrich) and according to (Dorrell et al, 2024). 24 DNAse-treated RNA libraries (4 conditions x 3 biological replica x 2 genetically distinct cell lines) were sequenced on a DNBseq Illumina platform (BGI Genomics Co., Ltd, Hongkong, China) with 100 bp paired-end sequencing. Raw reads were filtered by removing adaptor sequences, contamination and low-quality reads (reads containing over 40% bases with Q value < 20%) to obtain clean reads. Clean reads were mapped to the version 3 annotation of the *P. tricornutum* genome (Rastogi et al., 2018), and average TPM values for each gene in each library were calculated using DiatomicBase.

## Supporting information

Supplementary Informations

## Author contributions

AF and CB conceived and supervised the project. EV coordinated the project, provided initial databases and drafted the manuscript. NZ designed the website, loaded the data with the technical help of PV. SL designed and performed RNA-Seq analysis under progressive Fe limitation. HCC, RGD, CD, RM and AF performed and interpreted case studies. KV, MF, HCC, RGD, AF and CB provided critical suggestions to the manuscript. All authors proofread and approved the manuscript.

## Acknowledgements

The authors would like to thank the research community working on molecular aspects of diatoms for their feedback during the creation and curation of the website. NZ and EV thank Dr Ge and his team from the South Dakota State University for help in deploying iDEP on DiatOmicBase. NZ would like to thank Catherine Le Bihan, Maël Lefeuvre, Nolwenn Lavielle, Phi Phong Nguyen from the IBENS computing service for their technical support. We thank Mariella Ferrante, Svenja Mager, and Anna Santin for helping us to add *Pseudo-nitzschia multistriata* to DiatOmicBase. KV acknowledges Michiel Van Bel and Emmelien Vancaester for technical assistance and data curation within PLAZA Diatoms.

This work was supported principally by a grant from Gordon and Betty Moore Foundation GBMF8752. CB acknowledges additional support from the European Research Council (ERC) under the European Union’s Horizon 2020 research and innovation programme (Diatomic; grant agreement No. 835067), the French Government ‘Investissements d’Avenir’ programs MEMO LIFE (ANR-10-LABX-54) and PSL Research University (ANR-11-IDEX-0001-02). AF acknowledges funding from the Fondation Bettencourt-Schueller (Coups d’élan pour la recherche francaise-2018), the “Initiative d’Excellence” program (Grant “DYNAMO,” ANR-11-LABX-0011-01) and EMBRC-FR – “Investments d’avenir” program (ANR-10-INBS-02). AF and CB acknowledges the ANR BrownCut (Project-ANR-19-CE20-0020) and Horizon Europe BlueRemediomics (grant agreement No. 101082304) projects. HCC acknowledges funding from ANR DiaLincs (ANR 19-CE43-0011-01). RGD acknowledges an ERC Starting Grant (grant number 101039760, “ChloroMosaic”). KV acknowledges Research Foundation-Flanders (FWO) for ELIXIR Belgium [I002819N] and BOF/GOA No. 01G01715 and 01G01323. MF acknowledges a Villum Young Investigator Grant (Villum Fonden grant number 37521).

## Data statement

Raw fastq data corresponding to new RNA-Seq performed under progressive Fe limitation is provided in NCBI BioProject number PRJNA936812.

## Supporting information provided in a separate file

*Figure S1:* CPS III (Phatr3_J24195) genome browser capture displaying Phatr3 and Phatr2 annotations as well as peptides mapped using a proteogenomic pipeline. A zoom in the 5’ region shows that two peptide sequences are mapped on an exon predicted by Phatr3 gene model but not by Phatr2. This gene model is also predicted by mapping of RNA-Seq reads (here from a study of McCarthy et al., 2017).

*Figure S2* A snapshot of the complete gene page for CPS III (Phatr3_J24195).

*Figure S3* Heatmap showing two-component regulator gene expression in light- and nutrient stress-related transcriptomes.

*Figure S4* Heatmap displaying transcription factor expression in light and nutrient stress related experiments.

*Figure S5* Schematic diagram of the culture regime used for Fe limitation experiments ( BioProject number PRJNA936812).

