## Supplementary Informations for "DiatOmicBase, a gene-centered platform to mine functional omics data across diatom genomes"

### Supporting information

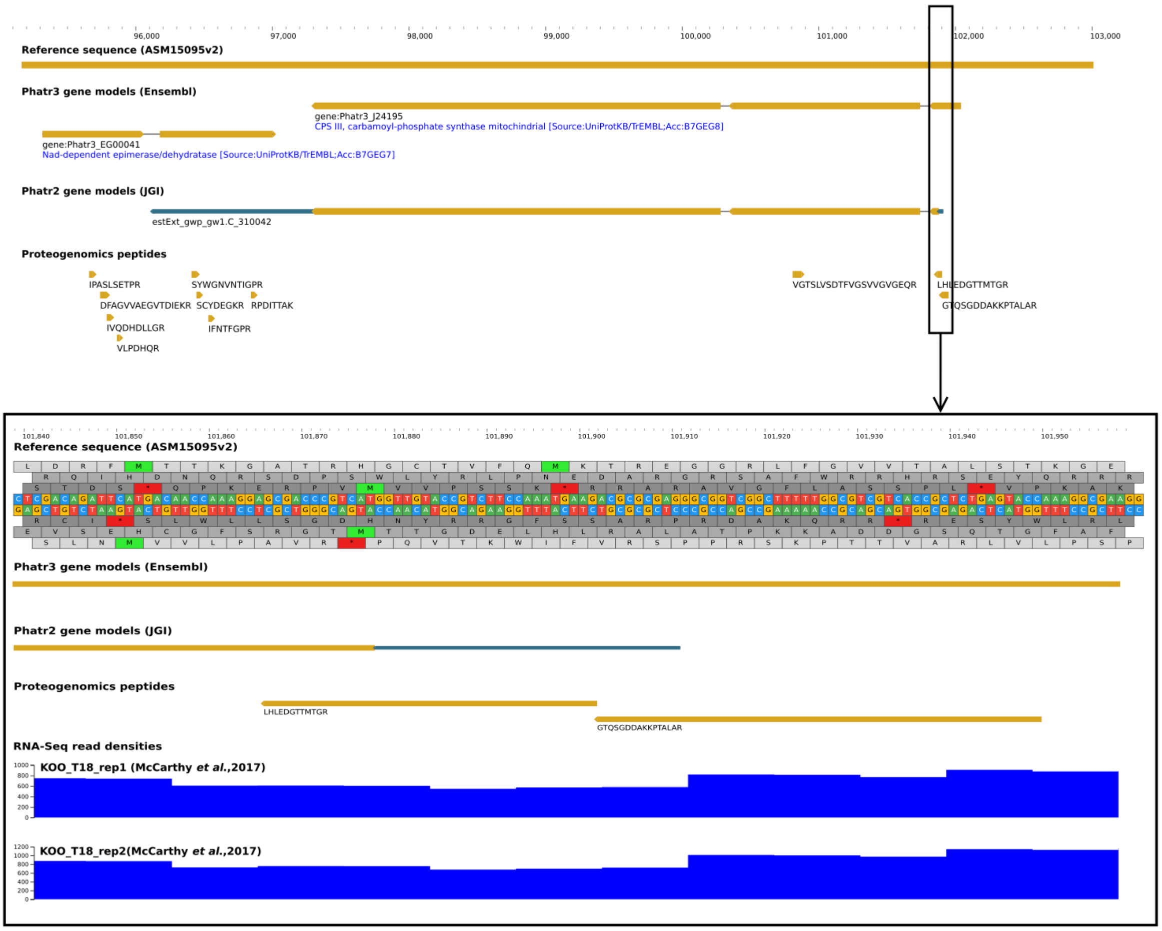


***Figure S1***

CPS III (Phatr3_J24195) genome browser capture displaying Phatr3 and Phatr2 annotations as well as peptides mapped using a proteogenomic pipeline. A zoom in the 5' region shows that two peptide sequences are mapped on an exon predicted by Phatr3 gene model but not by Phatr2. This gene model is also predicted by mapping of RNA-Seq reads (here from a study of McCarthy et al., 2017).

DiatOmicBase
All Species

Type your gene | All Species | Search

Home | Gene Page | Genome Browser | Resources | Transcriptomics | Help

### Gene: Phatr3_J24195

Download

Description: CPS III, carbamoyl-phosphate synthase mitochondrial [Source:UniProtKB/TrEMBL;Acc:B7GEG8]

Biotype: protein coding

Position: Chromosome 31:97309-102038

Additional ID:

- UniProtKB: B7GEG8
- NCBI: PHATRDRAFT_24195
- EMBL: CM000633
- Symbol: unCPS
- PLAZA: ptri02400
- Ensembl: Phatr3_J24195

| Phatr2 gene ID | Phatr2 protein ID | Phatr3 vs Phatr2 Category |
|---|---|---|
| estExt_gwp_gw1.C_310042 | 24195 | Modified |

### Sequence

### Genome Browser

Click on "**TRACKS SELECTOR**" (top left) to display additional information on the browser: Phatr2 prediction, ecotype variants, histone marks, ncRNAs...

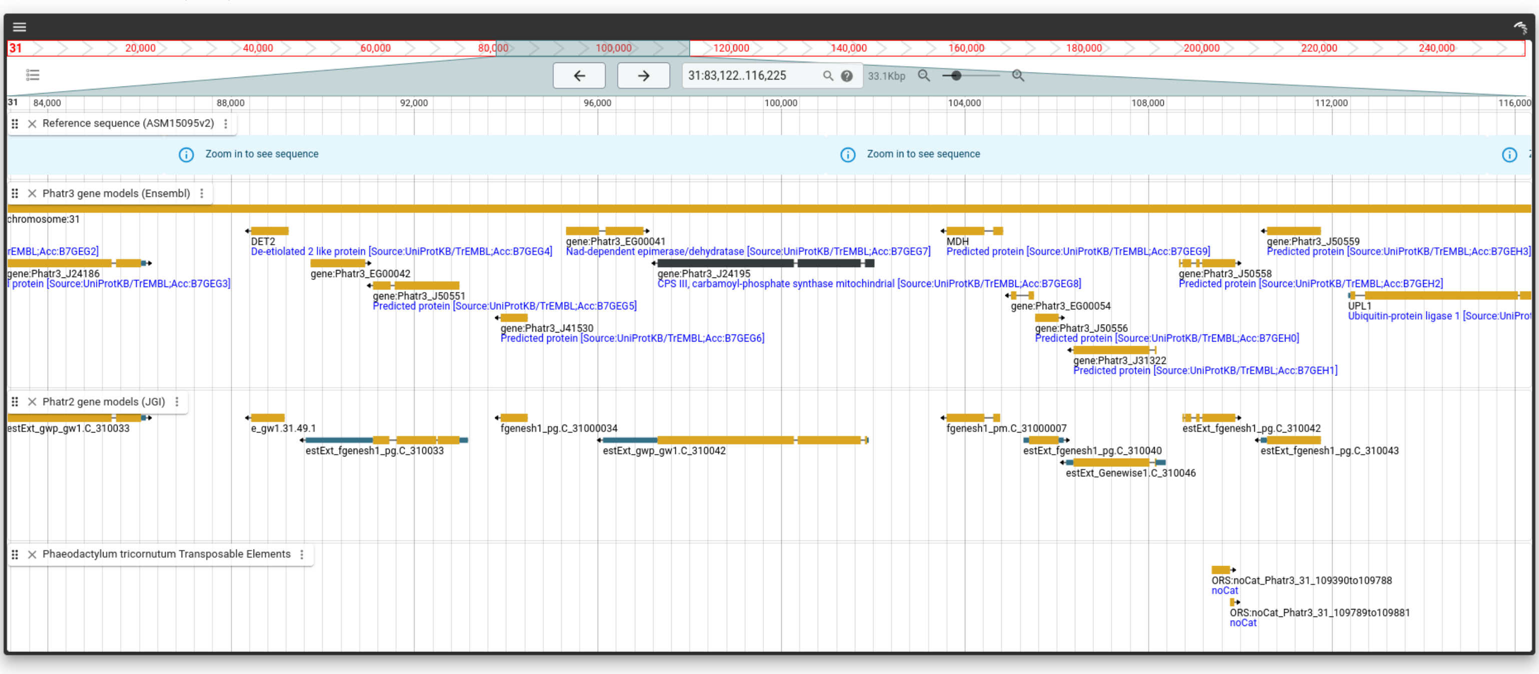


### Expression (RNA-Seq)

Transcripts were considered as differentially expressed if e-value < 0.05 with log2 fold change <-1 for down-regulated and log2FC >1 for up-regulated. Log2FC, adjusted e-value and experiment informations are available by clicking on the plot bars.

Download

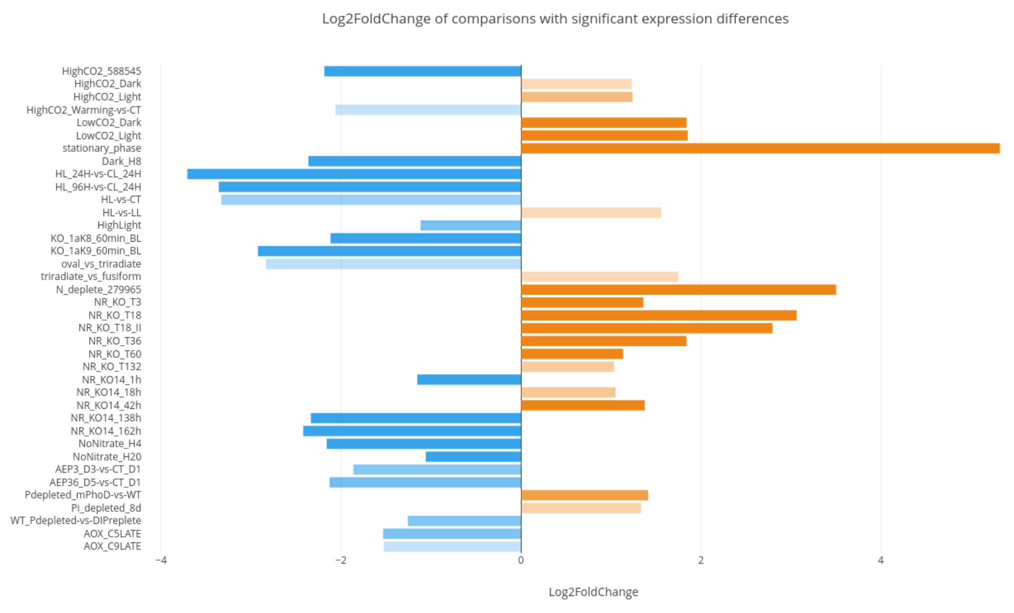


NON SIGNIFICANT

NON EXPRESSED

### GO terms

| | | |
|---|---|---|
| Molecular Function | GO:0005524 | ATP binding |
| Molecular Function | GO:0004088 | carbamoyl-phosphate synthase (glutamine-hydrolyzing) activity |
| Molecular Function | GO:0046872 | metal ion binding |
| Biological Process | GO:0006541 | glutamine metabolic process |
| Biological Process | GO:0006207 | 'de novo' pyrimidine nucleobase biosynthetic process |

### Domains

| | | |
|---|---|---|
| InterPro | IPR016185 | PreATP-grasp_dom_sf |
| InterPro | IPR036914 | MGS-like_dom_sf |
| InterPro | IPR011607 | MGS-like_dom |
| InterPro | IPR017926 | GATASE |
| InterPro | IPR035686 | CPSase_GATase1 |
| InterPro | IPR029062 | Class_I_gatase-like |
| InterPro | IPR005483 | CbamoylP_synth_lsu_CPSase_dom |
| InterPro | IPR005479 | CbamoylP_synth_lsu-like_ATP-bd |
| InterPro | IPR036480 | CarbP_synth_ssu_N_sf |
| InterPro | IPR002474 | CarbamoylP_synth_ssu_N |
| InterPro | IPR006274 | CarbamoylP_synth_ssu |
| InterPro | IPR036897 | CarbamoylP_synth_lsu_oligo_sf |
| InterPro | IPR005480 | CarbamoylP_synth_lsu_oligo |
| InterPro | IPR006275 | CarbamoylP_synth_lsu |
| InterPro | IPR013815 | ATP_grasp_subdomain_1 |
| InterPro | IPR011761 | ATP-grasp |
| CDD | cd01744 | GATase1_CPSase |
| Gene3D | 1.10.1030.10 | |
| Gene3D | 3.30.1490.20 | |
| Gene3D | 3.40.50.1380 | |
| Gene3D | 3.40.50.880 | |
| Gene3D | 3.50.30.20 | |
| HAMAP | MF_01209 | CPSase_S_chain |
| PRINTS | PR00098 | CPSASE |
| PROSITE | PS50975 | ATP_GRASP |
| PROSITE | PS00866 | CPSASE_1 |
| PROSITE | PS00867 | CPSASE_2 |
| PROSITE | PS51273 | GATASE_TYPE_1 |
| PROSITE | PS51855 | MGS |
| Pfam | PF02786 | CPSase_L_D2 |
| Pfam | PF02787 | CPSase_L_D3 |
| Pfam | PF00988 | CPSase_sm_chain |
| Pfam | PF00117 | GATase |
| Pfam | PF02142 | MGS |
| SMART | SM01096 | CPSase_L_D3 |
| SMART | SM01097 | CPSase_sm_chain |
| SMART | SM00851 | MGS |
| SUPFAM | SSF48108 | |
| SUPFAM | SSF52021 | |
| SUPFAM | SSF52317 | |
| SUPFAM | SSF52335 | |
| SUPFAM | SSF52440 | |
| TIGRFAMs | TIGR01369 | CPSaseII_lrg |
| TIGRFAMs | TIGR01368 | CPSaseIIsmall |

### KEGG

| | |
|---|---|
| K01948 | CPS1; carbamoyl-phosphate synthase (ammonia) [EC:6.3.4.16] |

### KOG

| | |
|---|---|
| KOG0370 | carbamoyl-phosphate synthase (glutamine-hydrolyzing) activity |

### Comparative Genomics

**Comparative genomics** data for this gene are hosted on the Diatoms PLAZA instance here.

*PLAZA is an access point for comparative and functional genomics centralizing genomic data produced by different genome sequencing initiatives. It integrates sequence data and comparative genomics methods and provides an online platform to perform evolutionary analyses and data mining.*

### Co-Expression Networks

Micro-Arrays data are available on DiatomPortal

#### Phaeonet Card:

**magenta**

Enriched in:

00910 Nitrogen metabolism [PATH:ko00910]

02000 Transporters [BR:ko02000]

### Other Supporting informations

**Associated Epigenomic Marks:**

- H3K27me3
- H3K9me2
- H3K9_14Ac
- H3K4me2

**Proteogenomics:**

| Phatr2 gene ID | Detection in Yang 2018 | Potential Non-Coding gene | Post translational modification |
|---|---|---|---|
| estExt_gwp_gw1.C_310042 | Yes | na | na |

**Histone methylation:**

☑ Polycomb marked gene

**Alternative splicing under N stress**

☐ Intron retention ☐ Exon skipping

**Targeting predictions:**

ASAfind Other
HECTAR mitochondrion
MitoFates Possessing mitochondrial presequence

**WolfPSort**

Animal: Mitochondria
Plant: Mitochondria
Fungi: Mitochondria
CONSENSUS: Mitochondria

| Evolutionary origins | Excluding ochrophytes and secondary plastids | Excluding SAR and CCTH |
|---|---|---|
| Lineage | SAR | Amorphea |
| Sub-group | Oomycetes | Opisthokonts |
| Number of consecutive top hits | 4 | 1 |
| Local diversity in top hits | No | No |

### Comments

Here you can post additional information about this gene: mutant, expression studies, gene model prediction... (reviewed data only).
To leave a comment, please specify: your name, your lab and the publication you refer to.

*Before posting, please make sure your comment follows the comments policy*

Login

Comment:
Name:
Lab:

ADD LABELS | ADD COMMENT

M↓ MARKDOWN HELP?

Upvotes Newest Oldest

**Emilie Villar**
0 points · 3 years ago
METABOLISM NITROGEN

Comment: This gene has been described as coding for a key enzyme of the ornithine-urea cycle by Allen et al. 2011.

Name: Emilie Villar

Lab: EV Consulting / IBENS

Publication: https://doi.org/10.1038/nature10074

**Emilie Villar**
0 points · 15 months ago

Comment: This gene has homology to the protein Thaps3a*40323 (THAPSDRAFT*40323) experimentally localized in the mitochondria.

Name:Emilie Villar

Lab:EV Consulting / IBENS

Publication:https://doi.org/10.1093/pcp/pcz097

Powered by **Commento++**

***Figure S2***

A snapshot of the complete gene page for CPS III (Phatr3_J24195)

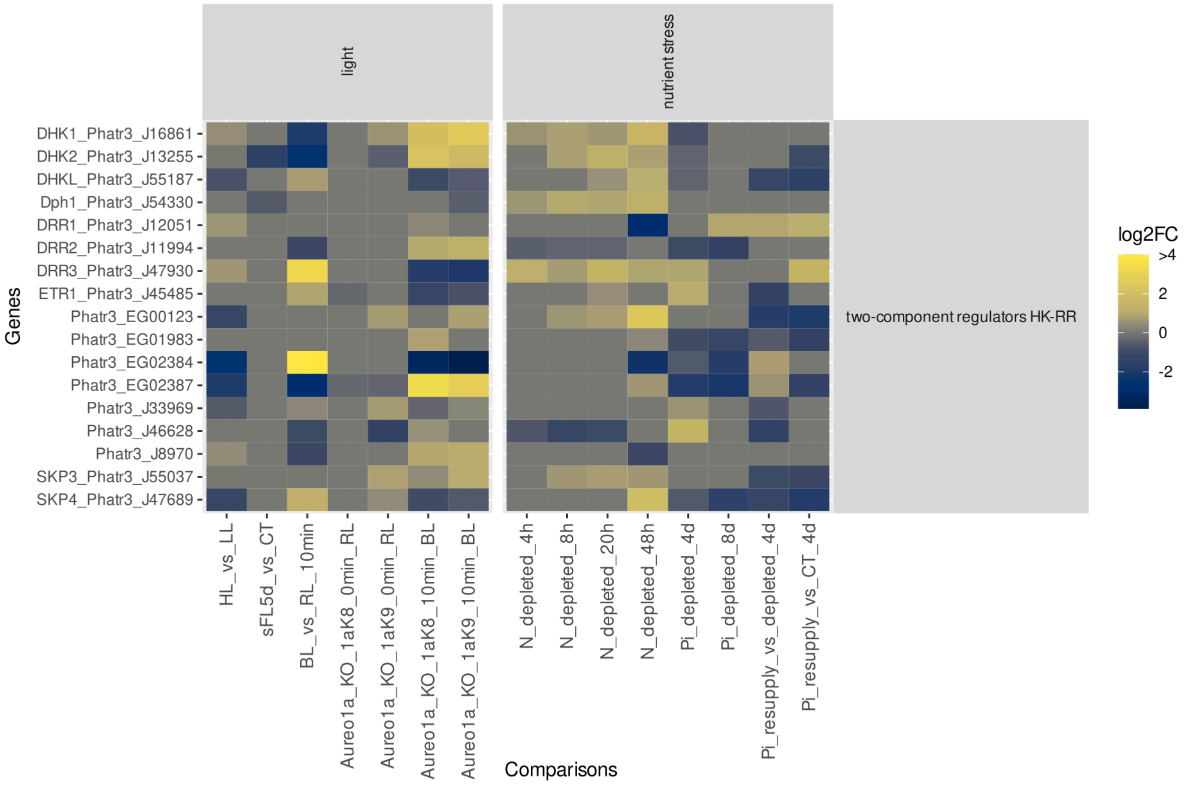


***Figure S3***

Heatmap showing two-component regulator gene expression in light- and nutrient stress-related transcriptomes.

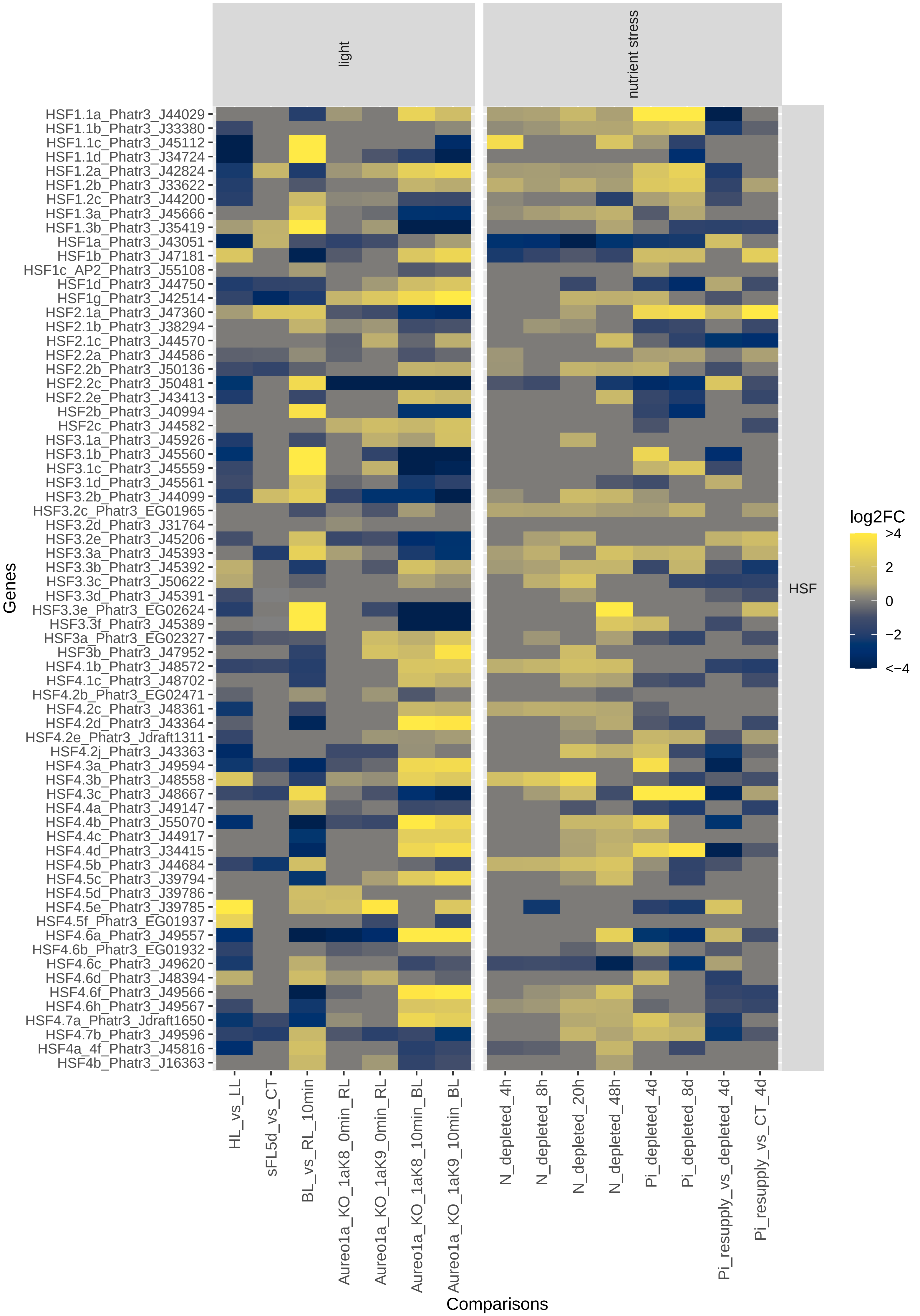
light
nutrient stress
HSF
Genes
HSF1.1a_Phatr3_J44029
HSF1.1b_Phatr3_J33380
HSF1.1c_Phatr3_J45112
HSF1.1d_Phatr3_J34724
HSF1.2a_Phatr3_J42824
HSF1.2b_Phatr3_J33622
HSF1.2c_Phatr3_J44200
HSF1.3a_Phatr3_J45666
HSF1.3b_Phatr3_J35419
HSF1a_Phatr3_J43051
HSF1b_Phatr3_J47181
HSF1c_AP2_Phatr3_J55108
HSF1d_Phatr3_J44750
HSF1g_Phatr3_J42514
HSF2.1a_Phatr3_J47360
HSF2.1b_Phatr3_J38294
HSF2.1c_Phatr3_J44570
HSF2.2a_Phatr3_J44586
HSF2.2b_Phatr3_J50136
HSF2.2c_Phatr3_J50481
HSF2.2e_Phatr3_J43413
HSF2b_Phatr3_J40994
HSF2c_Phatr3_J44582
HSF3.1a_Phatr3_J45926
HSF3.1b_Phatr3_J45560
HSF3.1c_Phatr3_J45559
HSF3.1d_Phatr3_J45561
HSF3.2b_Phatr3_J44099
HSF3.2c_Phatr3_EG01965
HSF3.2d_Phatr3_J31764
HSF3.2e_Phatr3_J45206
HSF3.3a_Phatr3_J45393
HSF3.3b_Phatr3_J45392
HSF3.3c_Phatr3_J50622
HSF3.3d_Phatr3_J45391
HSF3.3e_Phatr3_EG02624
HSF3.3f_Phatr3_J45389
HSF3a_Phatr3_EG02327
HSF3b_Phatr3_J47952
HSF4.1b_Phatr3_J48572
HSF4.1c_Phatr3_J48702
HSF4.2b_Phatr3_EG02471
HSF4.2c_Phatr3_J48361
HSF4.2d_Phatr3_J43364
HSF4.2e_Phatr3_Jdraft1311
HSF4.2j_Phatr3_J43363
HSF4.3a_Phatr3_J49594
HSF4.3b_Phatr3_J48558
HSF4.3c_Phatr3_J48667
HSF4.4a_Phatr3_J49147
HSF4.4b_Phatr3_J55070
HSF4.4c_Phatr3_J44917
HSF4.4d_Phatr3_J34415
HSF4.5b_Phatr3_J44684
HSF4.5c_Phatr3_J39794
HSF4.5d_Phatr3_J39786
HSF4.5e_Phatr3_J39785
HSF4.5f_Phatr3_EG01937
HSF4.6a_Phatr3_J49557
HSF4.6b_Phatr3_EG01932
HSF4.6c_Phatr3_J49620
HSF4.6d_Phatr3_J48394
HSF4.6f_Phatr3_J49566
HSF4.6h_Phatr3_J49567
HSF4.7a_Phatr3_Jdraft1650
HSF4.7b_Phatr3_J49596
HSF4a_4f_Phatr3_J45816
HSF4b_Phatr3_J16363
HL_vs_LL
sFL5d_vs_CT
BL_vs_RL_10min
Aureo1a_KO_1aK8_0min_RL
Aureo1a_KO_1aK9_0min_RL
Aureo1a_KO_1aK8_10min_BL
Aureo1a_KO_1aK9_10min_BL
N_depleted_4h
N_depleted_8h
N_depleted_20h
N_depleted_48h
Pi_depleted_4d
Pi_depleted_8d
Pi_resupply_vs_depleted_4d
Pi_resupply_vs_CT_4d
Comparisons
log2FC
>4
2
0
−2
<−4

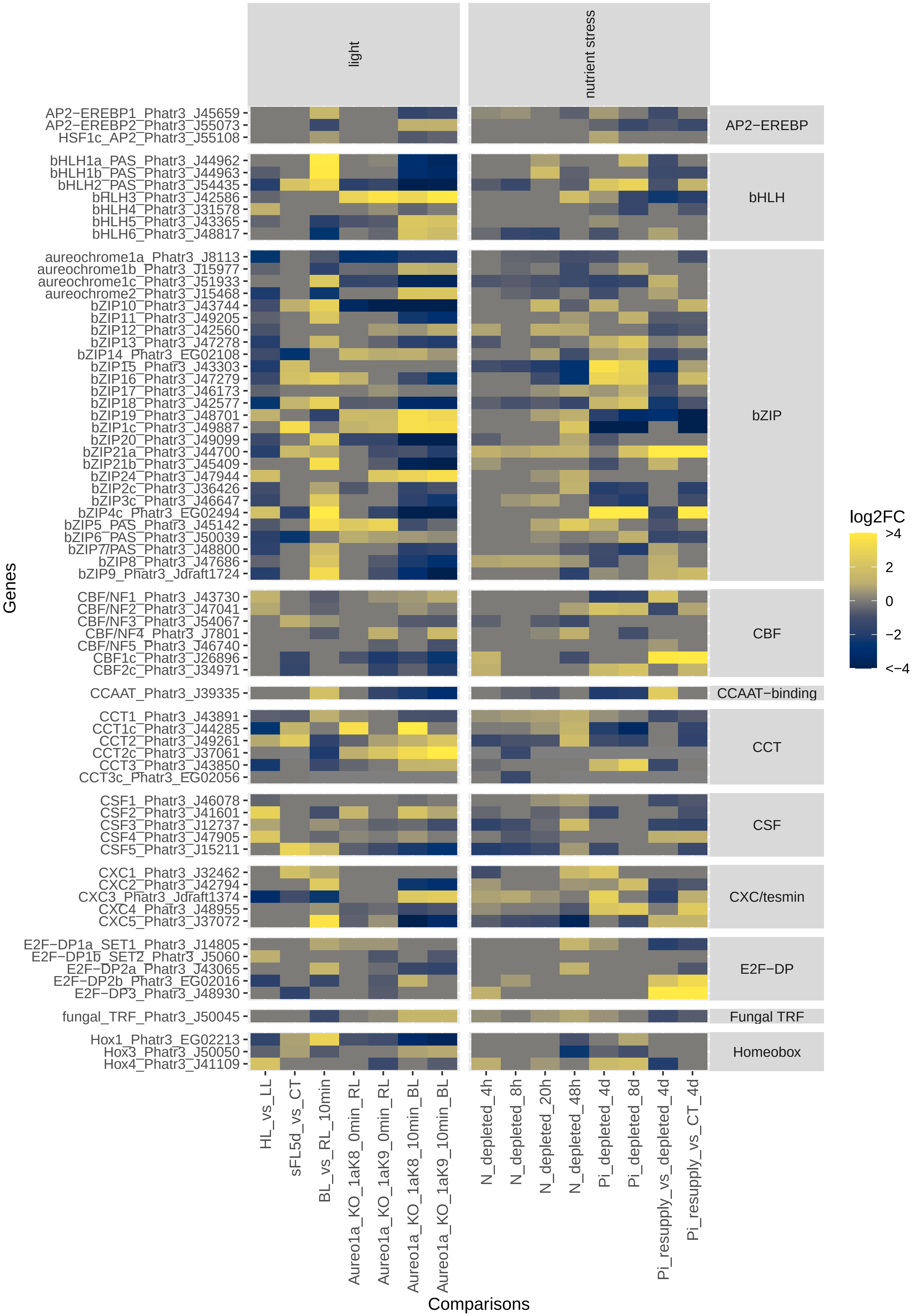
light
nutrient stress
AP2−EREBP1_Phatr3_J45659
AP2−EREBP2_Phatr3_J55073
HSF1c_AP2_Phatr3_J55108
bHLH1a_PAS_Phatr3_J44962
bHLH1b_PAS_Phatr3_J44963
bHLH2_PAS_Phatr3_J54435
bHLH3_Phatr3_J42586
bHLH4_Phatr3_J31578
bHLH5_Phatr3_J43365
bHLH6_Phatr3_J48817
aureochrome1a_Phatr3_J8113
aureochrome1b_Phatr3_J15977
aureochrome1c_Phatr3_J51933
aureochrome2_Phatr3_J15468
bZIP10_Phatr3_J43744
bZIP11_Phatr3_J49205
bZIP12_Phatr3_J42560
bZIP13_Phatr3_J47278
bZIP14_Phatr3_EG02108
bZIP15_Phatr3_J43303
bZIP16_Phatr3_J47279
bZIP17_Phatr3_J46173
bZIP18_Phatr3_J42577
bZIP19_Phatr3_J48701
bZIP1c_Phatr3_J49887
bZIP20_Phatr3_J49099
bZIP21a_Phatr3_J44700
bZIP21b_Phatr3_J45409
bZIP24_Phatr3_J47944
bZIP2c_Phatr3_J36426
bZIP3c_Phatr3_J46647
bZIP4c_Phatr3_EG02494
bZIP5_PAS_Phatr3_J45142
bZIP6_PAS_Phatr3_J50039
bZIP7/PAS_Phatr3_J48800
bZIP8_Phatr3_J47686
bZIP9_Phatr3_Jdraft1724
CBF/NF1_Phatr3_J43730
CBF/NF2_Phatr3_J47041
CBF/NF3_Phatr3_J54067
CBF/NF4_Phatr3_J7801
CBF/NF5_Phatr3_J46740
CBF1c_Phatr3_J26896
CBF2c_Phatr3_J34971
CCAAT_Phatr3_J39335
CCT1_Phatr3_J43891
CCT1c_Phatr3_J44285
CCT2_Phatr3_J49261
CCT2c_Phatr3_J37061
CCT3_Phatr3_J43850
CCT3c_Phatr3_EG02056
CSF1_Phatr3_J46078
CSF2_Phatr3_J41601
CSF3_Phatr3_J12737
CSF4_Phatr3_J47905
CSF5_Phatr3_J15211
CXC1_Phatr3_J32462
CXC2_Phatr3_J42794
CXC3_Phatr3_Jdraft1374
CXC4_Phatr3_J48955
CXC5_Phatr3_J37072
E2F−DP1a_SET1_Phatr3_J14805
E2F−DP1b_SET2_Phatr3_J5060
E2F−DP2a_Phatr3_J43065
E2F−DP2b_Phatr3_EG02016
E2F−DP3_Phatr3_J48930
fungal_TRF_Phatr3_J50045
Hox1_Phatr3_EG02213
Hox3_Phatr3_J50050
Hox4_Phatr3_J41109
AP2−EREBP
bHLH
bZIP
CBF
CCAAT−binding
CCT
CSF
CXC/tesmin
E2F−DP
Fungal TRF
Homeobox
Genes
HL_vs_LL
sFL5d_vs_CT
BL_vs_RL_10min
Aureo1a_KO_1aK8_0min_RL
Aureo1a_KO_1aK9_0min_RL
Aureo1a_KO_1aK8_10min_BL
Aureo1a_KO_1aK9_10min_BL
N_depleted_4h
N_depleted_8h
N_depleted_20h
N_depleted_48h
Pi_depleted_4d
Pi_depleted_8d
Pi_resupply_vs_depleted_4d
Pi_resupply_vs_CT_4d
Comparisons
log2FC
>4
2
0
−2
<−4

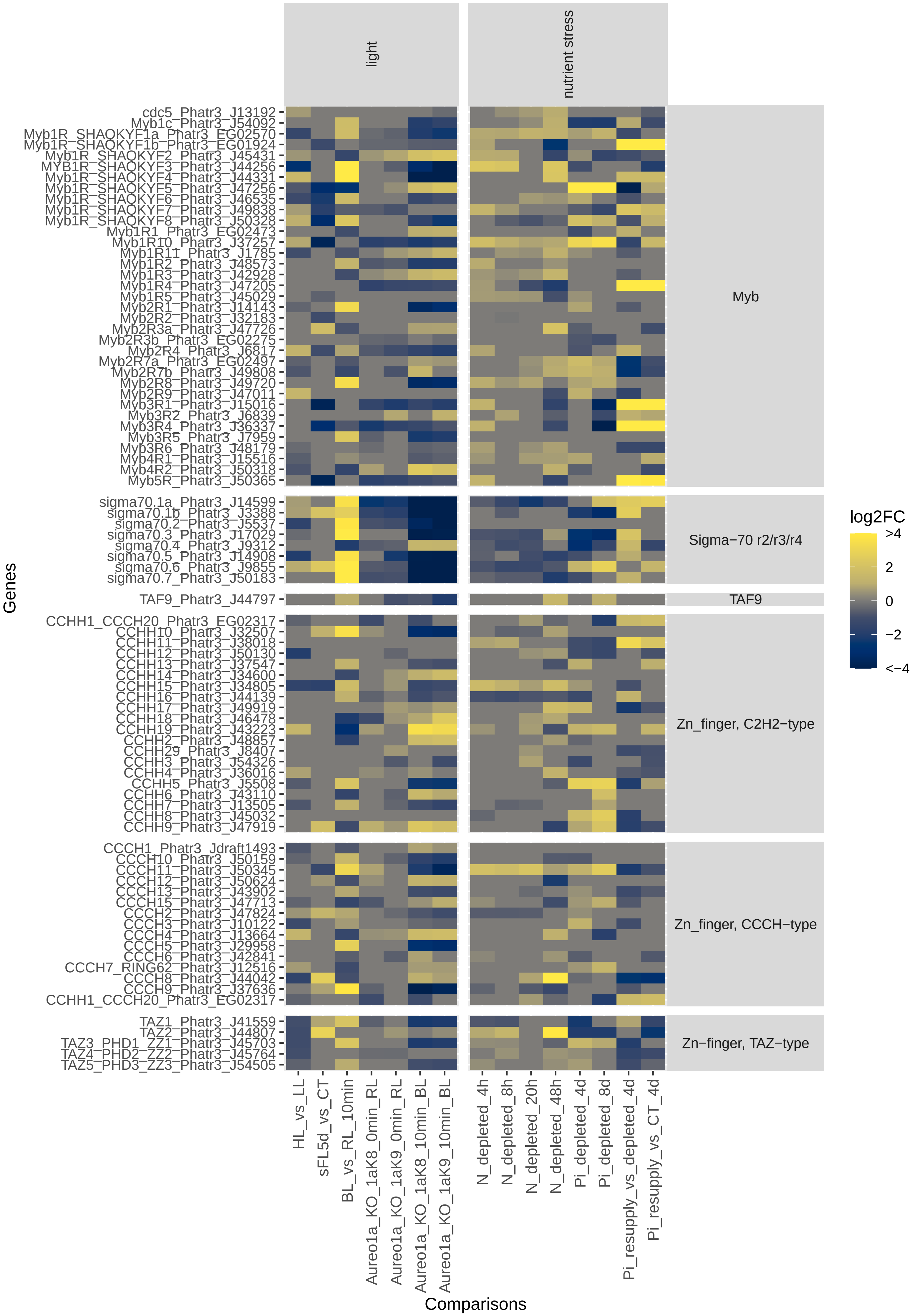
light
nutrient stress
cdc5_Phatr3_J13192
Myb1c_Phatr3_J54092
Myb1R_SHAQKYF1a_Phatr3_EG02570
Myb1R_SHAQKYF1b_Phatr3_EG01924
Myb1R_SHAQKYF2_Phatr3_J45431
MYB1R_SHAQKYF3_Phatr3_J44256
Myb1R_SHAQKYF4_Phatr3_J44331
Myb1R_SHAQKYF5_Phatr3_J47256
Myb1R_SHAQKYF6_Phatr3_J46535
Myb1R_SHAQKYF7_Phatr3_J49838
Myb1R_SHAQKYF8_Phatr3_J50328
Myb1R1_Phatr3_EG02473
Myb1R10_Phatr3_J37257
Myb1R11_Phatr3_J1785
Myb1R2_Phatr3_J48573
Myb1R3_Phatr3_J42928
Myb1R4_Phatr3_J47205
Myb1R5_Phatr3_J45029
Myb2R1_Phatr3_J14143
Myb2R2_Phatr3_J32183
Myb2R3a_Phatr3_J47726
Myb2R3b_Phatr3_EG02275
Myb2R4_Phatr3_J6817
Myb2R7a_Phatr3_EG02497
Myb2R7b_Phatr3_J49808
Myb2R8_Phatr3_J49720
Myb2R9_Phatr3_J47011
Myb3R1_Phatr3_J15016
Myb3R2_Phatr3_J6839
Myb3R4_Phatr3_J36337
Myb3R5_Phatr3_J7959
Myb3R6_Phatr3_J48179
Myb4R1_Phatr3_J15516
Myb4R2_Phatr3_J50318
Myb5R_Phatr3_J50365
Myb
sigma70.1a_Phatr3_J14599
sigma70.1b_Phatr3_J3388
sigma70.2_Phatr3_J5537
sigma70.3_Phatr3_J17029
sigma70.4_Phatr3_J9312
sigma70.5_Phatr3_J14908
sigma70.6_Phatr3_J9855
sigma70.7_Phatr3_J50183
Sigma−70 r2/r3/r4
TAF9_Phatr3_J44797
TAF9
CCHH1_CCCH20_Phatr3_EG02317
CCHH10_Phatr3_J32507
CCHH11_Phatr3_J38018
CCHH12_Phatr3_J50130
CCHH13_Phatr3_J37547
CCHH14_Phatr3_J34600
CCHH15_Phatr3_J34805
CCHH16_Phatr3_J44139
CCHH17_Phatr3_J49919
CCHH18_Phatr3_J46478
CCHH19_Phatr3_J43223
CCHH2_Phatr3_J48857
CCHH29_Phatr3_J8407
CCHH3_Phatr3_J54326
CCHH4_Phatr3_J36016
CCHH5_Phatr3_J5508
CCHH6_Phatr3_J43110
CCHH7_Phatr3_J13505
CCHH8_Phatr3_J45032
CCHH9_Phatr3_J47919
Zn_finger, C2H2−type
CCCH1_Phatr3_Jdraft1493
CCCH10_Phatr3_J50159
CCCH11_Phatr3_J50345
CCCH12_Phatr3_J50624
CCCH13_Phatr3_J43902
CCCH15_Phatr3_J47713
CCCH2_Phatr3_J47824
CCCH3_Phatr3_J10122
CCCH4_Phatr3_J13664
CCCH5_Phatr3_J29958
CCCH6_Phatr3_J42841
CCCH7_RING62_Phatr3_J12516
CCCH8_Phatr3_J44042
CCCH9_Phatr3_J37636
CCHH1_CCCH20_Phatr3_EG02317
Zn_finger, CCCH−type
TAZ1_Phatr3_J41559
TAZ2_Phatr3_J44807
TAZ3_PHD1_ZZ1_Phatr3_J45703
TAZ4_PHD2_ZZ2_Phatr3_J45764
TAZ5_PHD3_ZZ3_Phatr3_J54505
Zn−finger, TAZ−type
Genes
HL_vs_LL
sFL5d_vs_CT
BL_vs_RL_10min
Aureo1a_KO_1aK8_0min_RL
Aureo1a_KO_1aK9_0min_RL
Aureo1a_KO_1aK8_10min_BL
Aureo1a_KO_1aK9_10min_BL
N_depleted_4h
N_depleted_8h
N_depleted_20h
N_depleted_48h
Pi_depleted_4d
Pi_depleted_8d
Pi_resupply_vs_depleted_4d
Pi_resupply_vs_CT_4d
Comparisons
log2FC
>4
2
0
−2
<−4

***Figure S4***

Heatmap displaying transcription factor expression in light and nutrient stress related experiments.

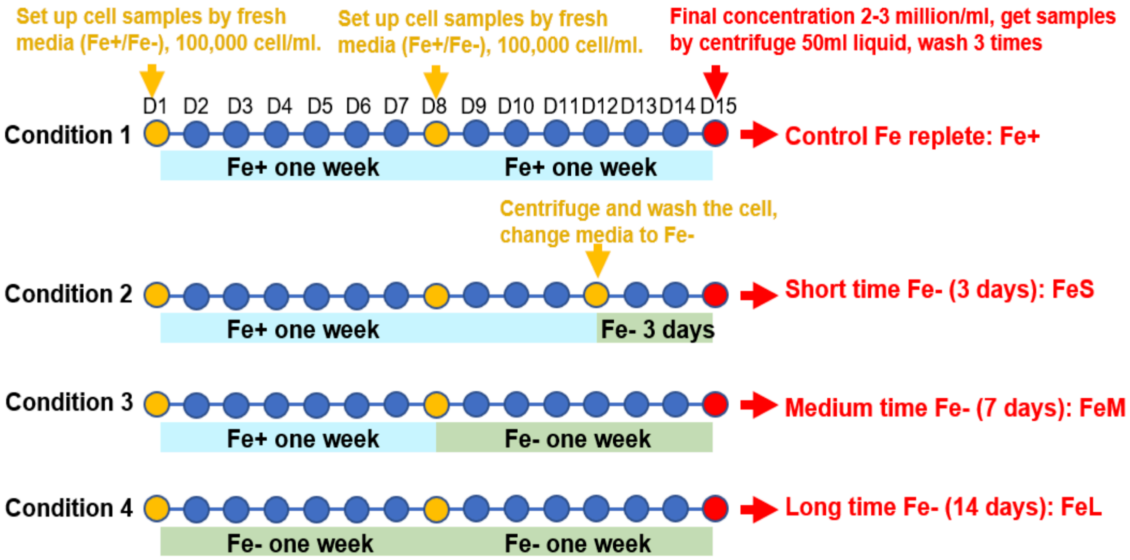


***Figure S5***

*Schematic diagram of the culture regime used for Fe limitation experiments ( BioProject number PRJNA936812).*
